# Potent Neutralization by Antibodies Targeting the Mpox A28 Protein

**DOI:** 10.1101/2025.05.22.655505

**Authors:** Ron Yefet, Leandro Battini, Mathieu Hubert, Katya Rakayev, Florence Guivel-Benhassine, Noam Rattner, Francoise Porrot, Lilach Abramovitz, Gilad Ostashinsky, Noam Ben-Shalom, Jeanne Postal, Ksenia Polonsky, Maya Ralph-Altman, Sireen Sweed, Tal Korner, Nadav Friedel, David Hagin, Eli Sprecher, Zvi Fishelson, Oren Kobiler, Lihi Adler-Abramovich, Olivier Schwartz, Pablo Guardado-Calvo, Natalia T Freund

## Abstract

Mpox is the most pathogenic Poxvirus in circulation. While several antigens have been identified as targets for neutralizing antibodies, many proteins remain unexplored. We isolated and characterized four monoclonal antibodies (mAbs) targeting the Mpox A28 (OPG153), a virulence factor present on mature Mpox virions. The antibodies were isolated from convalescent individuals, alongside 14 additional mAbs targeting the A35 and H3 proteins. Anti-A28 mAbs potently neutralized Mpox and Vaccinia virus (VACV) through complement-dependent mechanisms involving C1q and C3 deposition. High resolution crystal structures of Anti-A28 mAbs 10M2146 and 8M2110 in complex with VACV A26 revealed two proximal epitopes within the N-terminal domain. Passive transfer of 8M2110 attenuated disease in infected mice. Moreover, immunization with A28 elicited antigen-specific B cells and robust neutralizing antibody responses and provided complete protection against lethal VACV challenge. These findings support Mpox A28 as a promising target for the induction of neutralizing antibodies and antiviral interventions.

## INTRODUCTION

Mpox is a member of the Poxvirus family, an infamous viral lineage which has been a major source of human morbidity throughout history^1,2^. Ever since the successful eradication of Variola virus (the causative agent of Smallpox) in 1980, Mpox has emerged as the leading Poxvirus threat in circulation. Furthermore, cessation of mass vaccination campaigns has left most of the population increasingly vulnerable to Poxvirus infection^3,4^. Consequently, Mpox cases have steadily increased in endemic African countries, and eventually spread to non-endemic countries^1,4-6^. In 2022, the emergence of a new Mpox clade (clade IIb) led to swift dissemination of the virus to more than 100 non-endemic countries while infecting over 100,000 individuals^7^. To mitigate Mpox spread, an attenuated Vaccinia virus (VACV) based vaccine, Modified Vaccinia Ankara (MVA), was deployed to immunize at risk populations^8^. Although MVA confers protection from Mpox, new studies indicate a rapid decline of the elicited antibody response in the ensuing months, likely attributed to the attenuated nature and antigenic distance of MVA compared to Mpox^9-14^. This highlights the critical need to advance our understanding of the immune targets on Mpox, which is fundamental for developing new prevention and treatment strategies.

Mpox infection and vaccination elicits antibodies directed primarily against the surface proteins of its two infectious forms, Mature virions (MV) or Enveloped virions (EV), which are responsible for inter-host and intra-host transmission, respectively^15-22^. Out of the 30 different surface antigens expressed on the virus, six were so far described as targets for neutralizing monoclonal antibodies (mAbs). These include the MV attachment and fusion proteins A29, E8, H3, M1 and the EV proteins A35 and B6 (Mpox nomenclature)^21,23-26^. Some of these mAbs confer protection in animal models, most effectively when administered in a combination targeting both forms of the virus^21,27,28^. Moreover, the activity of human Poxvirus targeting mAbs is often complement-dependent^21,25^. The sheer abundance of viral targets recognized by antibodies elicited during infection, coupled with their largely unexplored mechanisms of viral neutralization, highlights the importance of identifying additional neutralizing antibodies targeting new viral sites and further exploring their mechanisms of action. Recent immunization studies in mice have indicated that additional Mpox surface proteins could serve as targets for neutralizing antibodies^29^. Among these, A28—a major structural component of the MV and a target of antibodies elicited by infection or vaccination—emerges as a promising candidate^22,30^. Acting as a virulence factor, A28 facilitates viral attachment through laminin binding and controls endosomal entry by regulating the viral fusion machinery^18,31,32^. While A28 represents a promising target for neutralizing antibodies, mAbs against this protein were not yet reported.

In this study, we characterize human monoclonal neutralizing antibodies targeting the Mpox A28 antigen. By analyzing B cells from Mpox convalescent donors, we generated a panel of mAbs against A28, as well as the previously described Mpox antigens, A35 and H3. Among these, mAbs targeting the A28 antigen demonstrated the highest neutralization, which was complement-dependent. Using X-ray crystallography, we obtained high-resolution structures of the two most effective anti-A28 mAbs, 10M2146 and 8M2110, in complex with the VACV A28 homolog, A26, showing that they bind two proximal, highly conserved, non-overlapping epitopes. The mode of action of both A28-targeting mAbs was elucidated using functional studies and immunogold transmission electron microscopy (TEM). Despite its potent in vitro neutralizing activity, passive administration of mAb 8M2110 in VACV-infected mice yielded only modest, non-significant clinical benefit in vivo. Nonetheless, A28 mice immunization elicited robust antigen-specific B cell responses and high-titer neutralizing antibodies, exhibiting both complement-dependent and -independent activities, and conferred complete protection against a lethal VACV challenge. Overall, our study presents A28 as a potential vaccine modality for elicitation of pan-Poxviruses neutralizing antibodies and provides functional and mechanistic insights about how human antibodies recognize and neutralize Mpox.

## RESULTS

### Generation of mAbs from human Mpox convalescent donors

Mpox infection elicits antibodies targeting both the MV and EV forms^15-17^. We previously reported that Mpox convalescent donors infected during the outbreak of May-June 2022 developed strong antibody and B cell responses against the A35 and H3 antigens^17^. Here, we isolated monoclonal antibodies (mAbs) from the same cohort while focusing on Mpox antigen A28 (OPG153), in addition to A35 (OPG 161) and H3 (OPG 108). To assess the serological response to A28, we recombinantly expressed residues 2-361 in a mammalian protein expression system (Figures 1A-1B) and tested serum reactivity of our cohort. A28 was strongly bound by 9 out of 11 convalescent sera (Figure S1A). Next, to isolate Mpox-specific B cells, we employed flow cytometry-based staining on selected convalescent donor samples collected 1-2-months post infection (indicated as V1) or 9-10-months post infection (indicated as V2). The peripheral blood mononuclear cells were stained with anti-CD19, anti-IgG and duo-labeled A28, as well as duo-labeled A35 and H3^17^ (Figure 1C). The identified cells were collected by single cell sorting, the mRNA of each B cell was extracted, and the membrane immunoglobulin heavy and light chains (Ig_H_ and Ig_L_, respectively) were amplified by PCR and Sanger-sequenced, as previously reported^33-36^. High-quality paired Ig_H_/Ig_L_ sequences were cloned and expressed as IgG1 mAbs. Four mAbs demonstrated binding to A28 (the mAbs 10M2146 and 10M2154, were clonal relatives), nine mAbs to A35 (the mAbs, 4M1130 and 4M1224, and the pair of mAbs, 4M1133 and 4M1166, were clonally related), and 5 mAbs to H3 (Figures 1D-1E).

**Figure 1:**
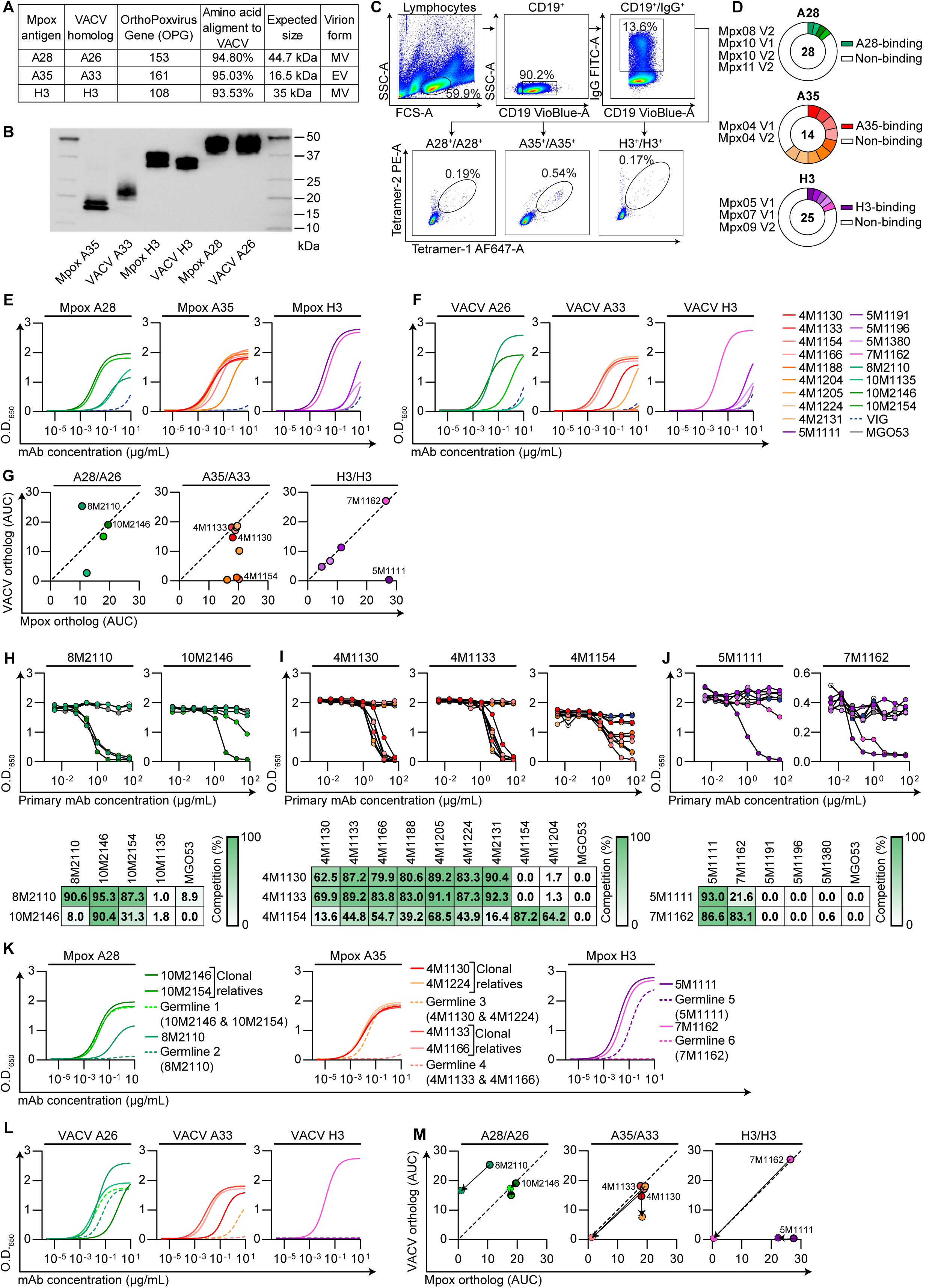
Generation of anti-A28, A33 and H3 Mpox binding mAbs. **(A)** Table listing the Mpox antigens and VACV homologs used in this study. **(B)** Western blot analysis using anti-Avi-tag HRP-conjugated secondary antibody of Mpox antigens and VACV homologs under reducing conditions. **(C)** Gating strategy for sorting enriched B cells from PBMCs of Mpox convalescent donors using the Mpox antigens listed in **A**, representative plots are shown. Lower left panel: using A28 (Mpx10 V2). Lower middle panel: using A35 (Mpx04 V1), Lower right panel: using H3 (Mpx05 V1). **(D)** Pie charts indicating the total IgG1 mAbs produced against each antigen (number listed in the middle). White slices represent non-binding mAbs. Colored slices represent individual mAbs that bound the corresponding antigen in ELISA. The Mpox convalescent donors from which the mAbs were isolated are indicated on the left. **(E)** Binding curves of anti-Mpox mAbs to their corresponding Mpox antigens A28, A35 and H3 as measured by ELISA of 12 consecutive 4-fold dilutions, starting from 10 μg/mL. Left panel: anti-A28 mAbs A. Middle panel: anti-A35 mAbs. Right panel: anti-H3 mAbs. **(F)** Same as **E** but for VACV homologs. **(G)** Differential two-dimensional representation of the AUC values from **E** and **F**. Dashed line represents line of identity. Left panel: for Mpox antigen A28 and the VACV A26 homolog. Middle panel: for Mpox antigen A35 and the VACV A33 homolog. Right panel: for Mpox antigen H3 and the VACV H3 homolog. **(H)** Degree of anti-A28 mAbs competition. Anti-A28 mAbs binding curves to Mpox A28 as measured by ELISA of 8 consecutive 4-fold dilutions, starting from 64 μg/mL. The anti-A28 mAbs competed against 0.2 μg/mL of the biotinylated 8M2110 (left) or 10M2146 (right). Table listing percentage of competition between mAbs is found on the bottom. **(I)** Same as **H** but for anti-A35 mAbs. The anti-A35 mAbs competed against 0.2 μg/mL of the biotinylated mAbs: 4M1130, 4M1133 and 4M1154 on the left, middle and right panels, respectively. **(J)** Same as **H** but for anti-H3 mAbs. The anti-H3 mAbs competed against 0.2 μg/mL of the biotinylated mAbs: 5M1111 and 7M1162 on the left and right panels, respectively. **(K)** Binding curves of germline reverted anti-Mpox mAbs to Mpox antigens A28, A35 and H3 as measured by ELISA of 12 consecutive 4-fold dilutions, starting from 10 μg/mL. Left panel: Anti-A28 mAbs. Middle panel: Anti-A35 mAbs. Right panel: Anti-H3 mAbs. **(L)** Same as **K** but for VACV homologs A26, A35 and H3. **(M)** Differential two-dimensional representation of the AUC values from **K** and **L**. Dashed line represents line of identity. Left panel: for Mpox antigen A28 and the VACV A26 homolog. Middle panel: for Mpox antigen A35 and the VACV A33 homolog. Right panel: for Mpox antigen H3 and the VACV H3 homolog. Arrows pointing from unmutated to germline reverted anti-Mpox mAb in dotted line. Anti-Mpox mAbs are depicted in the following color scheme: anti-A35 mAbs in shades of red, anti-H3 mAbs in shades of purple and anti-A28 mAbs in shades of green. Vaccinia immune globulin (VIG) is in dashed blue and MGO53 mAb serves as an isotype control is shown in gray. Germline reverted mAbs in **K-L** are in dashed lines in corresponding shading. Binding curves were determined by fitting values using Sigmoidal, 4PL (X is concentration) nonlinear regression for **E**, **F, K** and **L**.

We next tested whether the Mpox mAbs could also recognize their corresponding Vaccinia virus (VACV) homologs: A26 (A28, OPG 153), A33 (A35, OPG 161) and H3 (H3, OPG 108) all of which share a high sequence similarity (Figures 1A-1B)^37,38^. Most antibodies showed cross reactivity: three out of four A28 mAbs, six out of nine A35 mAbs and four out of five H3 mAbs, bound to both Mpox and VACV antigens. The remaining mAbs displayed preferential binding to either the Mpox or VACV orthologs, indicating recognition of species-specific domains (Figures 1E-1G). Competition ELISA pointed out that three of the four A28 mAbs (8M2110, 10M2146 and 10M2154) compete with one another, and therefore likely target proximal sites. Similarly, the nine A35 mAbs clustered into two main epitope groups, primarily based on their differential binding to the Mpox and VACV homologs. Two of the five H3 mAbs competed with each other, while the rest did not (Figures 1H-1J). This suggests that Mpox infection elicits antibodies directed against convergent immunodominant sites on A28, A35 and H3 antigens. These sites are partially conserved between VACV and Mpox antigen homologs, highlighting both shared and species-specific antigenic determinants.

The mAbs exhibited moderate levels of somatic hypermutations (SHMs). To explore whether affinity maturation plays a role in the antibody response to A28, A35, and H3, we selected nine mAbs representing the major antigenic clusters (based on competition ELISA) and reverted them to their predicted germline versions (indicated as ‘Germlines 1-6’, Figures 1K-1M). Predicted germlines 1-6 were generated based on V_H_D_H_J_H_ and V_L_J_L_ IMGT alignment, while keeping the CDRH3 intact as was previously reported^39-41^. Our data indicates that the binding of A28-targeting mAbs does not relay on SHM. As such, Germline 1, in which the somatic hypermutations of the clonally related anti-A28 mAbs 10M2146 (10 and 5 amino acid substitutions in Ig_H_ and Ig_L_, respectively) and 10M2154 (3 amino acid substitutions in Ig_H,_ none in Ig_L_) were reverted, demonstrated similar binding to either A28 or the VACV homolog. Germline 2—the germline-reverted 8M2110 (with 7 and 3 amino acid substitutions in Ig_H_ and Ig_L_, respectively)—lost its binding to A28, yet retained its binding to the VACV homolog A26 (Figures 1K–1M). These findings suggest that affinity maturation contributed mostly to the cross reactivity with other Poxviruses, while the ability to target the particular antigenic site is mostly germline-encoded. A mixed pattern was observed with mAbs targeting the A35 antigen. Germline 3, the unmutated precursor of mAbs 4M1130 and 4M1224 (each harboring only 2 amino acid substitutions in the heavy chain), showed only a slight reduction in binding compared to their mature counterparts. In contrast, Germline 4—corresponding to mAbs 4M1133 (with 6 and 1 amino acid substitutions in the heavy and light chains, respectively) and 4M1166 (7 substitutions in the heavy chain)—completely lost binding capacity (Figures 1K–1M). A similar trend was noted for anti-H3 mAbs: Germline 5, derived from mAb 5M1111 (5 and 3 substitutions in heavy and light chains), retained binding, while Germline 6 of mAb 7M1162 (also with 5 and 3 substitutions) showed a complete loss of reactivity. These findings suggest that while some Mpox-specific antibodies are germline-encoded, others require somatic hypermutation for effective binding.

### Antibodies targeting Mpox A28 exhibit potent complement-dependent neutralization

We next evaluated the neutralizing capacity of our mAbs against Mpox (clade IIb), as well as against three VACV strains, WR, Copenhagen and IHDJ. Several mAbs demonstrated neutralizing activity against Mpox or VACV, which was in all cases complement-dependent (except for the anti-A35 mAb 4M2131 that demonstrated a weak, but detectable, neutralization against VACV WR without complement, Figures 2A-2C). Complement-dependent neutralization is consistent with previous reports of human antibodies elicited following infection^17,21,25,42^. Among the tested mAbs, those targeting A28 exhibited the most potent neutralization. Their efficacy varied between viral species and across VACV strains. The clonal pair 10M2146 and 10M2154 neutralized Mpox with IC_50_ values of 0.65 μg/mL and 1.57 μg/mL, respectively, and the VACV IHDJ strain, with IC_50_ values of 2.61 μg/mL and 72.58 μg/mL, respectively. MAb 8M2110 neutralized Mpox with IC_50_ value of 9.47 μg/mL and displayed IC_50_ values of 0.06 μg/mL and 0.16 μg/mL against VACV WR and IHDJ, respectively (Figure 2C). VACV Copenhagen was resistant to all tested anti-A28 mAbs, likely a result of the absence of the A26 antigen on the viral surface due to a truncation of its C-terminal region^43-45^. The ability of anti-A28 mAbs to neutralize both the VACV WR and IHDJ strains highlights the potential of mAbs directed against A28 to exhibit broad pan-Poxvirus neutralization. To our knowledge, this is the first report to describe human Mpox-neutralizing mAbs targeting the A28 protein or its homologs. Among the anti-H3 mAbs, 7M1162 was the only one to neutralize all three tested VACV strains, with IC_50_ values of 1.18 μg/mL, 8.85 μg/mL, and 16.93 μg/mL against WR, Copenhagen, and IHDJ, respectively (Figures 2B-2C). However, it did not neutralize Mpox. In contrast, the Mpox-specific mAb 5M1111 neutralized Mpox with an IC_50_ of 39.17 μg/mL but had no activity against VACV. In this assay, anti-A35 mAbs exhibited minimal detectable neutralization activity (Figures 2A-2B).

**Figure 2:**
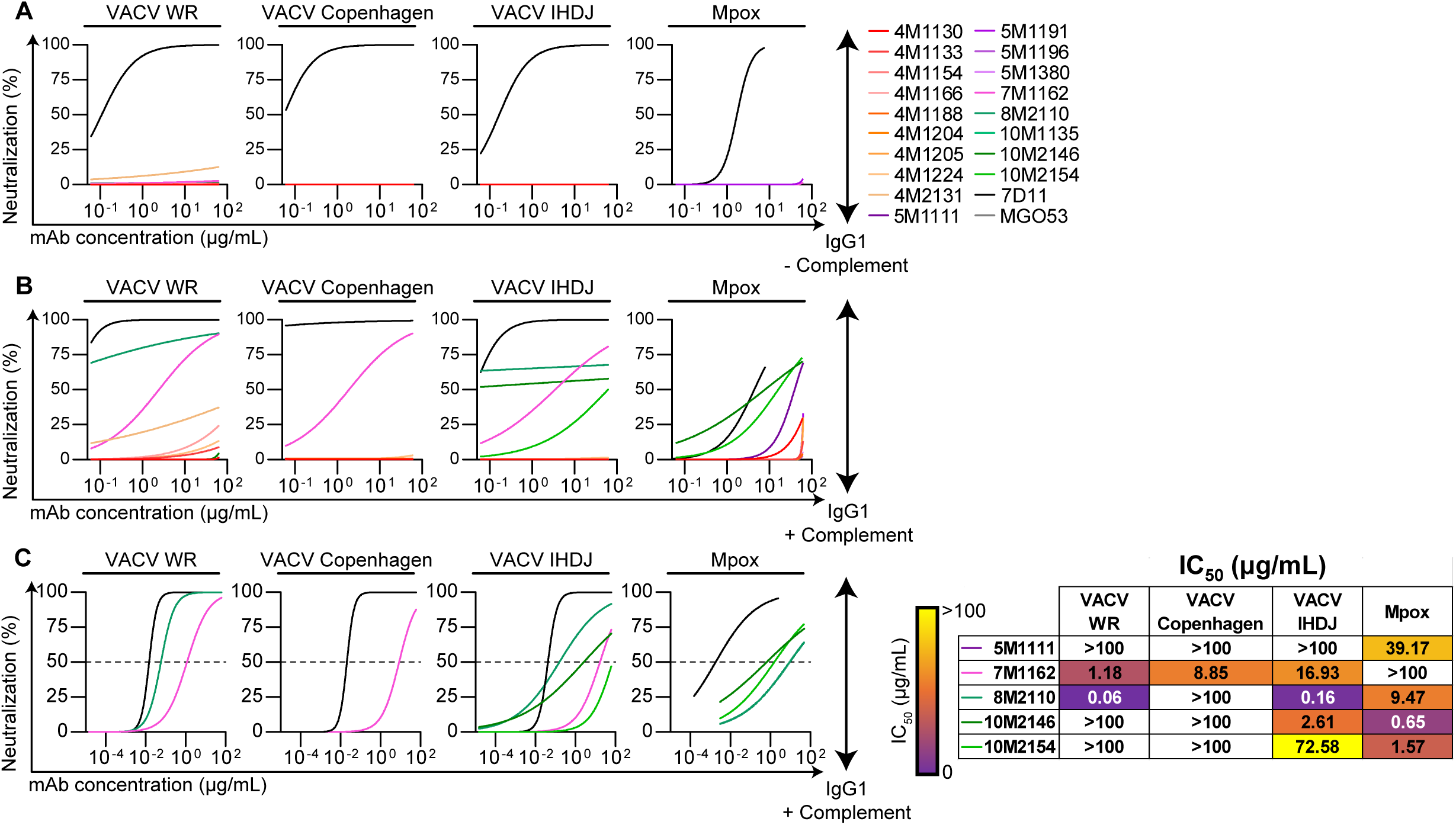
Anti-Mpox mAbs neutralization is complement dependent. **(A)** Neutralization of anti-Mpox mAbs against VACV strains and Mpox in the absence of complement. Neutralization curves of 6 consecutive 4-fold dilutions, starting from 62.5 μg/mL. From left to right, panels represent neutralization of VACV WR, VACV Copenhagen, VACV IHDJ strains and Mpox. **(B)** Same as **A** but in the presence of 1.5% NHS or 10% GPC, for VACV strains and Mpox, respectively. **(C)** Neutralization of anti-Mpox mAbs against Mpox and VACV strains in the presence of 1.5% NHS. Neutralization curves of 12 consecutive 4-fold dilutions, starting from 62.5 μg/mL for VACV strains and 8 consecutive 4-fold dilutions, starting from 50 μg/mL for Mpox. Dashed line represents f 50% neutralization. From left to right, panels represent neutralization of VACV WR, VACV Copenhagen, VACV IHDJ strains and Mpox. A heatmap with the corresponding IC_50_ values is on the right. Anti-Mpox mAbs are depicted in the following color scheme: anti-A35 mAbs in shades of red, anti-H3 mAbs in shades of purple and anti-A28 mAbs in shades of green. Anti-M1 mAb 7D11(black) is used as a positive control, while MGO53 (gray) is used as an isotype control. Neutralization curves and IC_50_ values were determined by fitting values using the Agonist vs. normalized response (Variable slopes) nonlinear regression for **A**-**C**.

### MAb combinations targeting different surface proteins enhance Mpox neutralization

Simultaneous targeting of multiple antigens on the surface of viruses is advantageous^46^, and was previously shown to enhance Poxvirus neutralization in vitro and in animal models^21,27,28^. To evaluate the efficacy of different mAb combinations, we tested our 18 mAbs in pairs. For Mpox, as well VACV strain IHDJ, combinations targeting 2 different antigens exhibited significantly stronger neutralization compared to combinations targeting a single antigen (Figures 3A-3D). This was not the case for VACV strains WR and Copenhagen. When the two most potent mAbs targeting different antigens were tested alone and in combination across 12 serial dilutions, nearly all combinations showed a significant reduction in IC_50_ and IC_80_ values compared to single mAbs (Figures 3E and S2A-S2C). Notably, a synergistic effect was observed between A28- and H3-targeting mAbs and anti-A35 mAbs, even though the latter exhibited minimal neutralization on their own.

**Figure 3:**
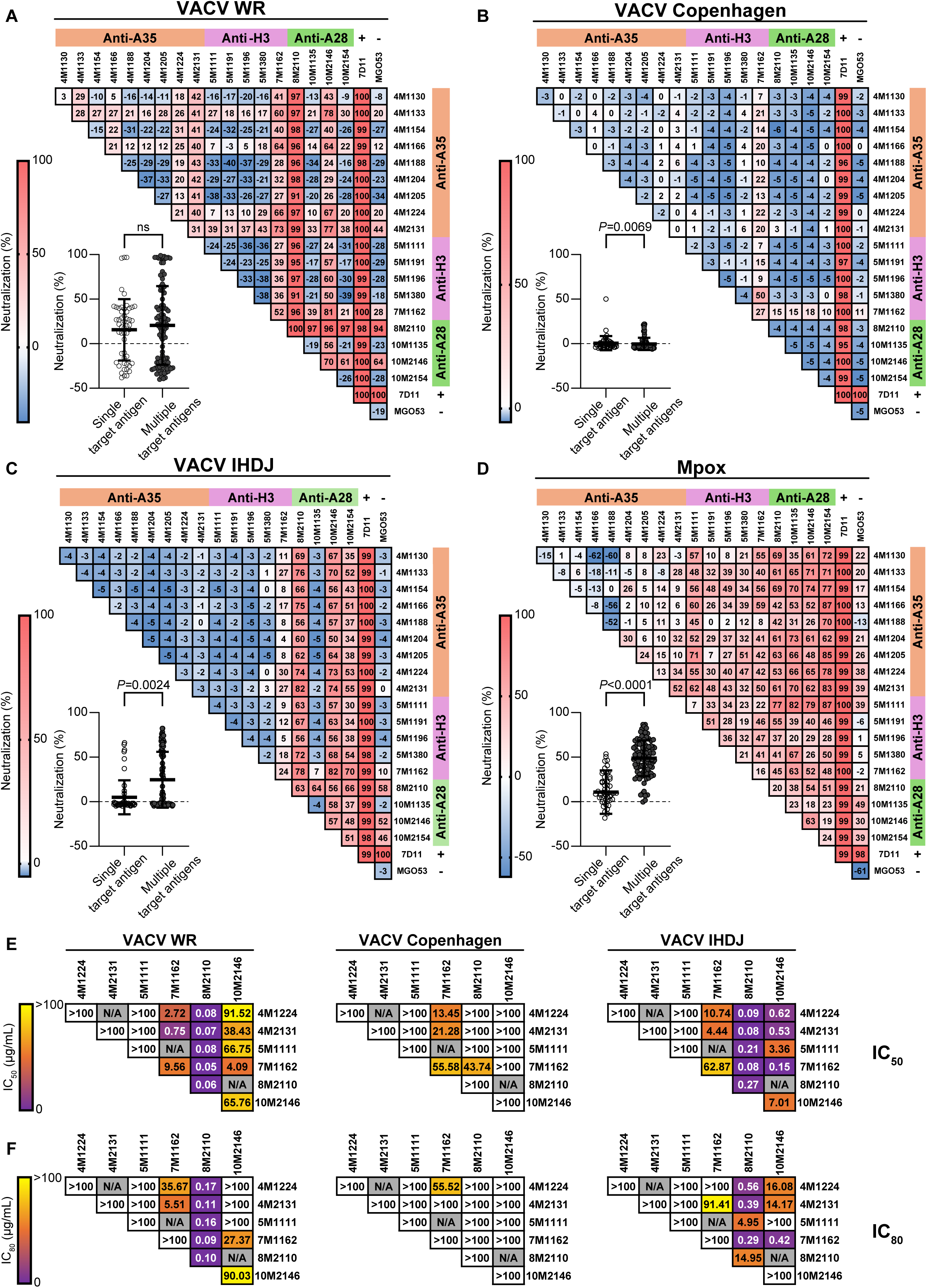
Poxvirus neutralization by anti-Mpox mAb combinations. **(A)** Heatmap demonstrating the neutralization values of VACV WR using anti-Mpox mAbs combinations in the presence of 1.5% NHS at a total mAb concentration of 62.5 μg/mL. Each mAb combination as normalized to baseline infection (no mAb). Neutralization Scale bar is found on the left. Lower left panel: analysis of neutralization values for anti-Mpox mAbs combinations (excluding 7D11 and MGO53) targeting single or multiple Mpox antigens. **(B)** Same as **A** but for VACV Copenhagen. **(C)** Same as **A** but for VACV IHDJ. **(D)** Same as **A** but for Mpox in the presence of 10% GPC. **(E)** Heatmap depicting summarized IC_50_ and IC_80_ values of select two-mAb combinations targeting two antigens, as shown in **Figure S3**. From left to right, panels represent VACV WR, VACV Copenhagen and VACV IHDJ strains. Scale bar is found on the left. For **A**-**D** mAbs combinations targeting a single Mpox antigen are in clear dots and mAbs combinations targeting multiple Mpox antigens are in dark gray dots. Statistical analysis was performed using Mann-Whitney test. Negative neutralization results represent combinations with higher infection level then the selected baseline infection well. ns= non significant.

### 8M2110 and 10M2146 mAbs bind two proximal epitopes of the N-terminal domain on A28

To further characterize anti-A28 neutralizing mAbs, we performed structural analyses focusing on the two most potent variates, members of two distinct clones, 8M2110 and 10M2146, each isolated from a different donor. For these studies, we utilized the VACV homolog of A28, A26. Structurally, A26 comprises two principal domains: the “head domain” (residues 1–320), which contains both the acid-sensing region (residues 1–75) and the fusion suppressor region (residues 76–320), both implicated in the pH-dependent entry of the virion; and the “tail domain” (residues 320–500), which includes the A27-binding region that anchors A26 to the viral membrane through its interaction with VACV A27 (homologous to Mpox A29)^47^. The head domain has been previously resolved and subdivided into a N-terminal domain (NTD; residues 1–228) and a C-terminal domain (CTD; residues 229–364) (Figure S3)^31^. The NTD consists of twelve α-helices (α1–α12) and contains two histidine residues (H48 and H53), which are essential for viral infectivity. The CTD adopts a mixed α/β fold composed of six α-helices (α13–α18) and six β-strands (β1–β6)^31^. For structural determination, both mAbs were expressed as F(ab) fragments and complexed with the residues 1-397 of A26, encompassing the full fusion-regulatory region.

We obtained the crystal structures of 8M2110:A26 and 10M2146:A26 at 2.8 Å and 1.9 Å resolution, respectively. In both complexes, A26 resembles its unbound conformation (rmsd = 0.2 Å), with no significant antibody-induced conformational changes. The paratope of 10M2146 is comprised mainly of the complementarity-determining regions (CDRs) H2, H3, L2, and L3, and the N-terminus of the light chain, and spans a buried surface area of 971 Å^2^ on the NTD, mainly involving helixes α7 and α8, with some minor contacts on α4 and α11 (Figures 4A-4B). Sequence analysis (Figure S3) indicates that two epitope residues differ in Mpox (F153 to L153, E156 to D156), but structural data suggest these substitutions are compatible with antibody binding. E156, which interacts with K78 and Y122 in the heavy chain, can be replaced by D156, and F153 can be replaced by L153 without introducing steric clashes (Figure 4B). The paratope of 8M2110 is formed mainly by CDRs H1, H2, and H3, with little to no contribution from the light chain. The epitope is spanning over 843 Å^2^, mostly (90%) on the NTD, involving residues in helixes α10-13 (Figures 4C-4D). Sequence analysis of the epitope revealed two residues that differ between VACV A26 and Mpox A28: A207 (V207 in Mpox) and F235 (L235 in Mpox) (Figure S3). Substituting A207 with V207 is expected to introduce some steric clashes with CDR-H1, which may explain why 8M2110 binds VACV A26 better than Mpox A28 (Figures 1E-1G and 4D). Mapping the footprints of both antibodies on A26 (Figure 4E) reveals that they bind two distinct epitopes that are very close to one another with some overlap on helix α11, explaining the competitive binding observed in ELISA assays (Figure 1H).

**Figure 4:**
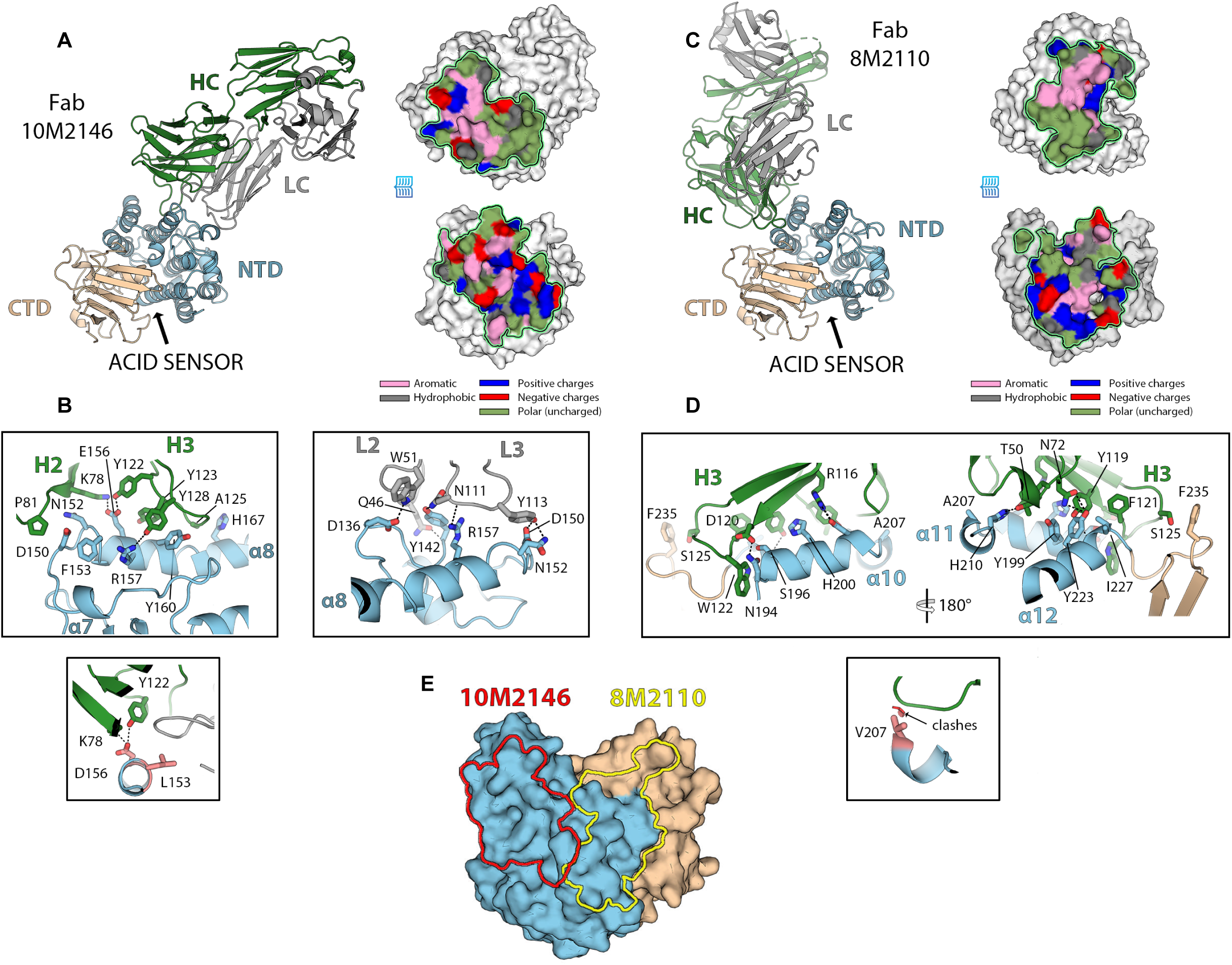
Crystal structures of anti-A28 neutralizing mAbs, 10M2146 and 8M2110. **(A)** Structure of the VACV A26/Fab-10M2146 complex as diffracted at 1.9 Å resolution. Left panel: The VACV A26 NTD and CTD are shown in blue and tan, respectively. The heavy chain of the Fab is colored green, the light chain in gray. An arrow marks the approximate location of the pH-sensing histidines. Right panel: “open book” views of the paratope (top) and epitope (bottom) outlined with a green line and colored based on the chemical nature of the residues, as indicated. **(B)** Close-up views of the 10M2146 epitope with the side chains involved in the interaction showed in sticks and labelled. Upper panels: interactions involving CDR H2 and H3 (left) and interactions involving CDR L2 and L3 (right). Lower panel: display of the two epitope residues that differ between Mpox and VACV, colored red and labelled, which have no impact on the interaction with the antibody. **(C)** Structure of the VACV A26/Fab-8M2110 complex, as diffracted at 2.8 Å resolution colored as in **A**. **(D)** Upper panel: Two close-up views of the 8M2110 epitope with the CDR H3 side chains involved in the interaction showed in sticks and labelled, as in **B**. Lower panel: displays the epitope residue V207 present in Mpox, which clashes with the antibody main chain. **(E)** Outline of the footprints of 10M2146 (red) and 8M2110 (yellow) on VACV A26.

The binding epitope of the anti-H3 mAb 7M1162 was predicted using 11 affinity selected peptides isolated from screening random phage display peptide libraries and computational prediction^48-51^, and an unbiased modeling analysis using AlphaFold3^52^. Both approaches resulted in different predicted clusters (1–3), with a single overlapping residue: histidine at position 167 (Figures S4A-S4C, S4E). Point mutants H3_H167G_, H3_H167K_ exhibited reduced to complete loss of binding to 7M1162, thus confirming this residue as part of the antibody binding site (Figure S4D). Notably, the H167K or H167G substitution did not affect binding of 5M1111 (Figure S4D).

### Anti-A28 mAbs block viral infection by recruiting C1q and C3

Consistent with previous reports on anti-Poxvirus antibodies^17,21,25,42,53,54^, most of the mAbs isolated in this study, required complement to effectively neutralize either VACV or Mpox. Moreover, heat inactivation at 56°C of the normal human serum (NHS) used as a source of complement abolished neutralization (Figure S5A). We therefore sought to decipher the mechanism of complement-dependent neutralization. For these assays we used VACV IHDJ, and the neutralizing mAbs 7M1162 (anti-H3) and 8M2110, 10M2146 and 10M2154. The strongest neutralizing effect was observed when antibodies were pre-incubated with the virus and 1.5% NHS for two hours before infection. Shortening this pre-incubation time drastically reduced antibody neutralization, resulting in 40% - 100% reduction when mAbs and NHS were added to U2OS cells at the time of infection (Figure 5A). These results suggest that the antibodies primarily act on the virus itself, rather than on infected cells, and that their neutralization potency builds up over time. Supporting this, anti-Mpox mAbs—despite binding to infected cells—did not trigger complement-dependent cytotoxicity (Figures S5B and S5C). We next investigated whether the mAbs promote complement-dependent virolysis as previously described^55^. However, inhibition of C5 using C5-IN-1 had no impact on neutralizing activity of our mAbs, suggesting that formation of membrane attack complex is unlikely to be the primary mechanism of antibody-mediated neutralization (Figure 5B). In contrast, blocking C3 with the selective inhibitor AMY-101 (CP40) significantly reduced neutralizing activity (Figure 5C).

**Figure 5:**
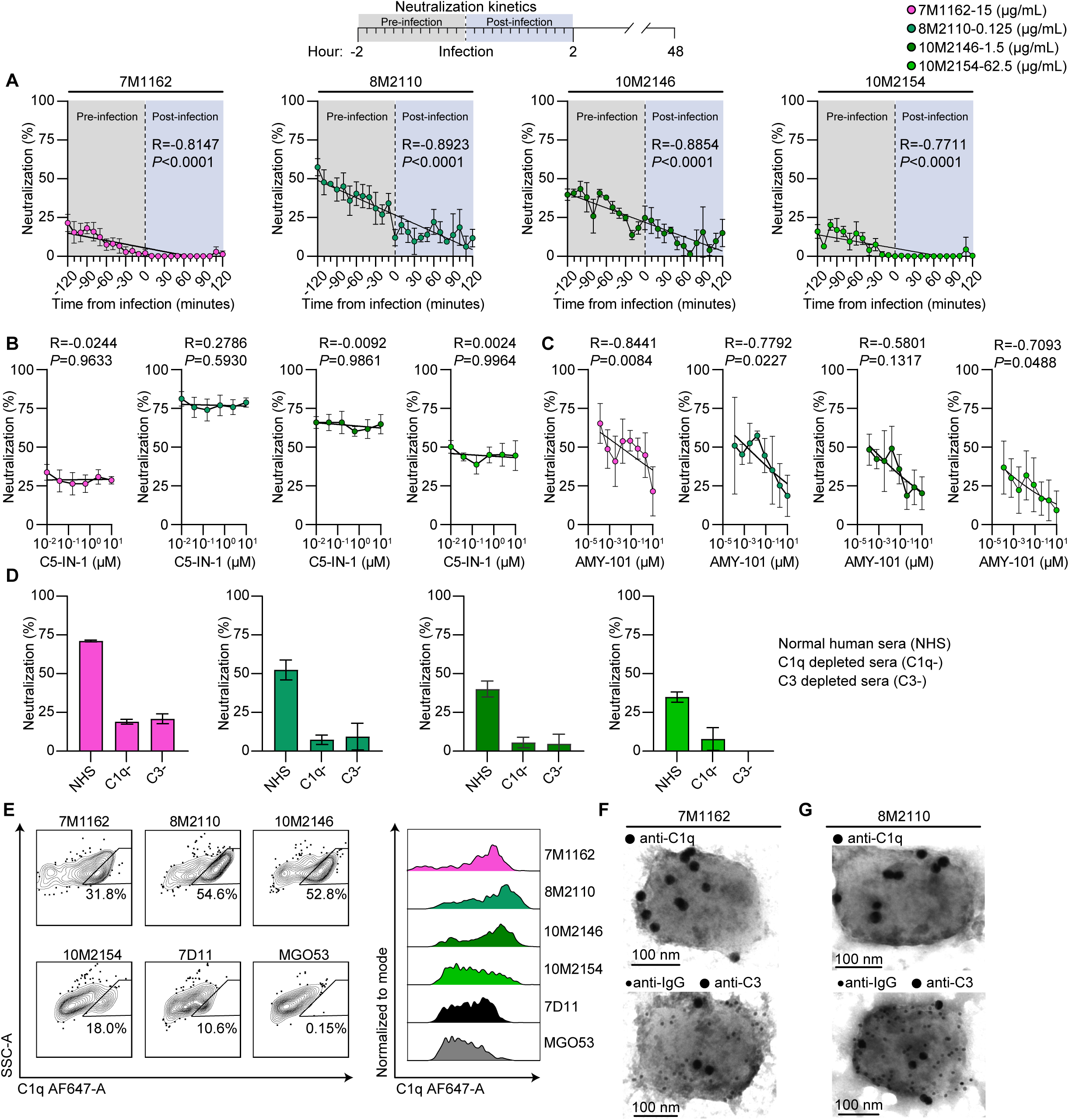
Anti-Mpox mAbs mechanism of complement dependent neutralization. **(A)** Neutralization kinetics of VACV IHDJ in the presence of anti-Mpox mAbs in IC_50_ concentrations and 1.5% NHS. NHS and mAbs were added in 10 minutes intervals from 2 hours pre-infection (mixed with the virus itself) to 2 hours post-infection. **(B)** Neutralization of VACV IHDJ in the presence of anti-Mpox mAbs in IC_50_ concentrations,1.5% NHS and 6 consecutive 4-fold dilutions, starting from 10 μM of a selective complement C5 inhibitor, C5-IN-1 (Compound 7). **(C)** Same as **B** but with 8 consecutive 5-fold dilutions, starting from 10 μM of a selective complement C3 inhibitor, AMY-101 (CP40). **(D)** Neutralization of VACV IHDJ in the presence of anti-Mpox mAbs in IC_50_ concentrations, 1.5% NHS or C1q-, or C3-depleted human serum. **(E)** C1q deposition on the membrane of VACV IHDJ virions. MV particles (∼5×10^6^ PFU) were incubated in the presence of 1.5% NHS and 10 μg/mL of listed mAbs. Left panel: flow cytometry plots, pre-gated for virions using GFP followed by staining for C1q-AF647. Right panel: Median florescent intensity. **(F)** TEM analysis of mAb deposition and complement factors C1q and C3 on VACV IHDJ virions. Purified MV particles were incubated with 10 μg/mL of mAb 7M1162 and 1.5% NHS. IgG, C1q, and C3 deposition were visualized using anti-human IgG gold nanoparticles (10 nm) or secondary mouse anti-C1q and mouse anti-C3, followed by labeling with anti-mouse gold nanoparticles (20 nm). **(G)** Same as F but for mAb 8M2110. Anti-Mpox mAbs are depicted in the following color scheme: anti-H3 mAb is in magenta and anti-A28 mAbs are in shades of green. For the neutralization assays shown in panels **A**–**D**, the following mAb concentrations, corresponding to their IC_50_ values, were used: 15 μg/mL for 7M1162, 0.125 μg/mL for 8M2110, 1.25 μg/mL for 10M2146, and 62.5 μg/mL for 10M2154. Spearman correlation was used in **A** to determine the relationship between pre-incubation times and neutralization. Standard curves were determined by fitting values using simple linear regression for **A** and Inhibitor vs. normalized response (Variable slopes) nonlinear regression for **B**-**C**. Anti-M1 7D11 (black) serves as a positive control and MGO53 (gray) serves as an isotype control. Standard deviation of mean for each value is shown in **A**– **D**.

The mAbs effectively bound the viral surface and promoted C1q deposition, however, exogenous C1q alone was insufficient for antibody-mediated neutralizing activity, suggesting that both C1q and C3 are essential for viral neutralization (Figures 5E and S5D–S5E). Serum depletion of either C1q or C3 impaired neutralization efficacy (Figure 5D), further strengthening our conclusion that mAb-mediated neutralization is facilitated by deposition of both C3 and C1q. TEM imaging confirmed the presence of IgG, C1q, and C3 on the viral surface (Figures 5F–5G), supporting a previously described mechanism named ‘full occupancy model’^53^, in which mAbs and C3 act as molecular ‘handcuffs,’ to immobilize the virus and prevent cellular entry.

### Antibodies targeting A28 mediate protection from lethal viral challenge in vivo

Prophylactic administration of 8M2110 to BALB/c mice (Figure 6A) did not confer protection from a lethal VACV challenge. Nevertheless, treated mice displayed a modest, although not statistically significant, delay in the onset of severe disease (weight *P*=0.056, survival *P*=0.1013), along with a slight reduction in viral load on day 5 post-infection compared to mice treated with isotype control antibody *P*=0.0883 (Figures 6B–6D). This trend was reproducible across three independent experiments. The gap between the potent in vitro neutralization capacity of 8M2110 and its limited in vivo efficacy is likely the result of the dominant EV form in intra-host viral dissemination, where the A28 antigen is not exposed.

**Figure 6:**
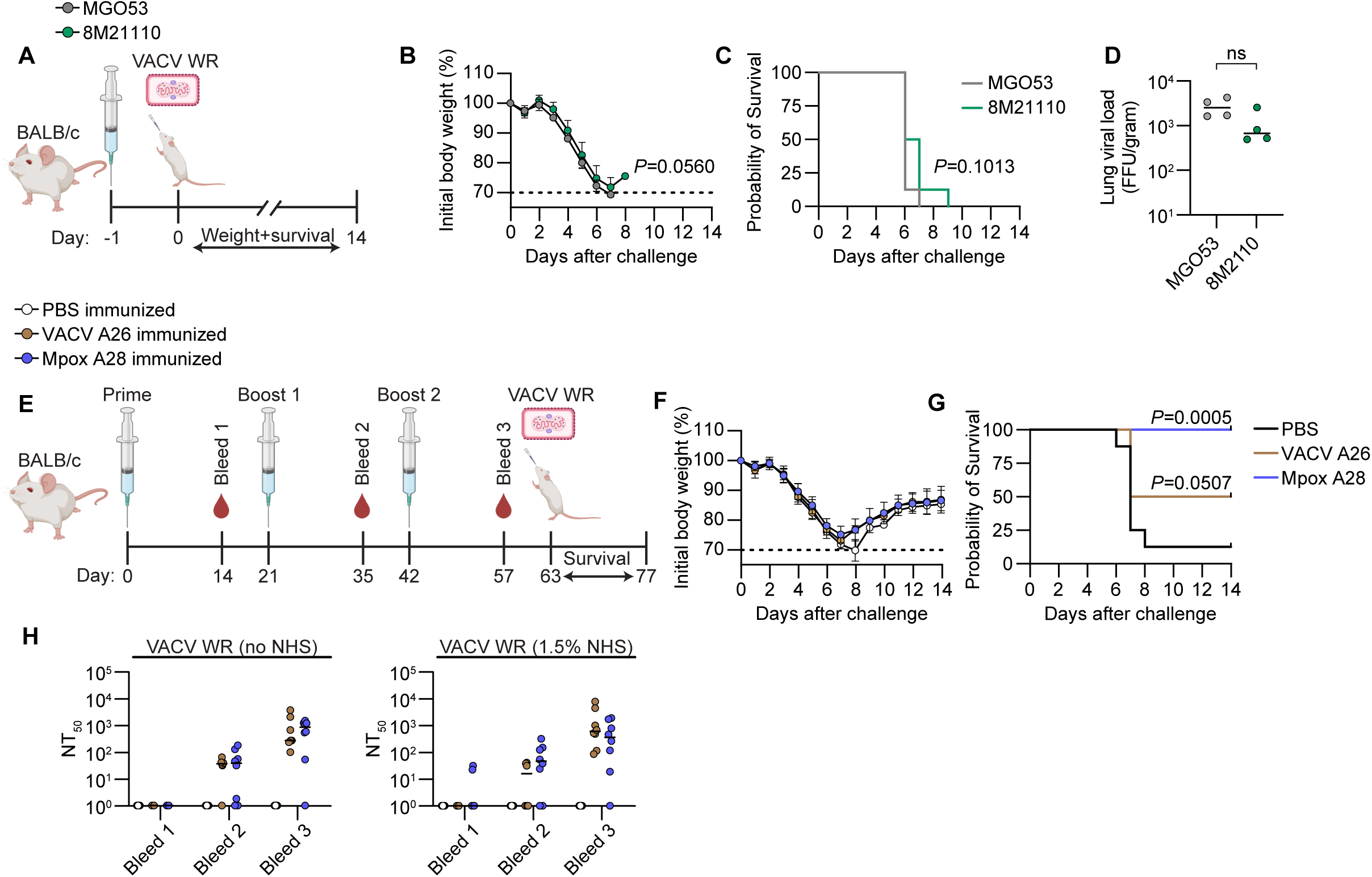
Antibodies against A28 confer protection from a lethal VACV challenge. **(A)** Schematic representation of mice prophylactic mAb transfer and challenge. BALB/c mice (n = 8 per group) were immunized I.P. with 200 μg of mAbs 8M2110 (anti-A28) or MGO53 (isotype control). The following day, mice were inoculated I.N. with a lethal dose of VACV WR (2 × 10^5^ PFU) and their weight and survival were monitored for 14 days. Mice that lost more than 30% of their initial weight (or more than 25% and exhibited a core body temperature below 34°C) were considered to have reached the no recovery threshold and were subsequently sacrificed. **(B)** Body weight changes of initial weight for 8M2110 or MGO53 immunized mice. **(C)** Survival of 8M2110 or MGO53 immunized mice. **(D)** Lung viral load on day 5 post-VACV challenge in BALB/c 8M2110 or MGO53 immunized mice (n = 4 per group). **(E)** Schematic representation of mice A28 immunization and challenge. BALB/c mice (n = 8 per group) were immunized I.P. with either PBS, Mpox A28, or VACV A26 antigens (10 μg per dose), formulated with QS-21 adjuvant. Immunizations were administered three times (prime + two boosts) at 3-week intervals. Two weeks after each injection, mice were bled and serum was collected and heat inactivated. Three weeks after the final boost, mice were challenged I.N. with a lethal dose of VACV WR (2 × 10^5^ PFU). Weight and survival were monitored daily for 14 days. Mice that lost more than 30% of their initial weight (or more than 25% and exhibited a core body temperature below 34°C) were considered to have reached the no recovery threshold and were subsequently sacrificed. **(F)** Body weight changes of initial weight for A28-, A26- or PBS-immunized mice. **(G)** Survival of for A28-, A26- or PBS-immunized mice. **(H)** Time course analysis of heat-inactivated sera neutralization of VACV WR strain with or without 1.5% NHS. NT_50_ values were calculated from neutralization curves obtained through six consecutive 4-fold dilutions, starting at 1:40. Left panel: neutralization in the absence of NHS. Right panel: neutralization in the presence of 1.5% NHS. Mice immunized with mAb 8M2110 are represented in green, while those immunized with MGO53 are shown in gray. PBS-immunized mice are in white, VACV A26-immunized mice are in brown, and Mpox A28-immunized mice are in blue. Body weight comparison analysis was performed using Main-effects analysis for **B**. In panels **C** and **H**, statistical analysis was conducted by comparing survival curves to the isotype control using the Log-rank (Mantel-Cox) test, with Holm-Sidak multiple comparison correction applied in **C** and by Unpaired T test in **D**. Images were created using BioRender. ns= non significant.

To assess whether vaccination with A28 elicits neutralizing antibodies capable of conferring protection, we immunized mice three times (both BALB/c and C57BL/6, see Methods) with either PBS, Mpox A28, or VACV A26 antigens formulated with QS-21 (saponin) or Alum, respectively (Figures 6E and S6B). Immunization with Mpox A28 conferred complete protection against a lethal VACV challenge (Figures 6F-6G and S6A). To assess whether this protection is mediated by antibodies we analyzed mouse sera and B cells. As expected, antigen-specific antibody titers increased following each immunization (Figure S6C). Furthermore, sera from vaccinated mice effectively neutralized both VACV WR and IHDJ strains (as anticipated, VACV Copenhagen was not neutralized, Figure S6D). Surprisingly, heat-inactivated sera retained neutralizing activity almost completely even in the absence of complement, suggesting that antibodies elicited following A28 immunization block viral attachment or entry through additional, complement-independent mechanisms (Figure 6E). Further characterization of the immune cell response of immunized mice revealed elevated levels of germinal center (GC) B cells and an increased frequency of A28-binding IgG^+^ B cells (Figures S6E-S6F), while no significant differences were observed in the total B cell (B220+), CD4+, or CD8+ T cell populations between the immunized groups (Figures S6E-S6F Notably, immunization with VACV A26 provided less protection than Mpox A28. Both BALB/c and C57BL/6 mice showed lower neutralizing antibody titers following A26 vaccination. Among the four A26-vaccinated mice that did not survive, neutralizing antibody levels were particularly low within their group. Together, these findings demonstrate that immunization with Mpox A28 induces a robust humoral immune response, marked by the production of neutralizing antibodies that act through both complement-dependent and -independent mechanisms, ultimately conferring effective protection against lethal Poxvirus challenge.

## DISCUSSION

Mpox outbreaks have become increasingly frequent in recent years, highlighting the need for novel targets for antibody-based interventions. In this study, we identify A28 protein as a dominant target of human neutralizing antibodies. Although A28 is not essential for infection, it critically regulates the transition from EV production, associated with intra-host spread, to MV production, which facilitates inter-host transmission^56^. Therefore, A28 contributes to viral dissemination and inter-individual transmission. The mAbs isolated from convalescent donors exhibited high-affinity binding to Mpox A28 and its VACV homolog A26 and demonstrated potent neutralization of Mpox and multiple VACV strains. Structural analysis revealed that the neutralizing mAbs bind to distinct, proximal epitopes on the NTD of A28, providing insight into conserved immunogenic regions, and supporting A28 classification as a promising candidate for broad Poxvirus vaccine development. The identification of A28 as a potent target for neutralizing antibodies against Mpox advances our understanding of immune responses to Poxviruses and informs future therapeutic and vaccine strategies.

Anti-viral neutralizing mAbs often neutralize by blocking viral entry into host cells^57^. However, unlike viruses such as HIV-1, SARS-CoV-2, and influenza, where antibodies alone can effectively prevent infection^33,58-60^, Poxvirus neutralization frequently depends on complement factors. This complement-dependent mechanism, long recognized as a key feature of Poxvirus immunity^21,42,53,54^, was consistently observed among the potent anti-A28 and anti-H3 mAbs characterized here. Our mechanistic data demonstrates that antibody-mediated viral neutralization is facilitated by the deposition of complement components C1q and C3 on the viral surface. These complement factors likely enhance antibody effectiveness by overcoming the structural complexity of the Poxvirus entry machinery. Given the large size of Poxviruses and multiple pathways for host cell entry, complement deposition likely serves as an amplifier, working with antibodies to ensure efficient blockade of infection. TEM analysis further provided compelling visual confirmation of this “molecular handcuff” model, complying with the full occupancy model, in which antibodies immobilize virions via complement engagement, thereby preventing infection^53,54,57^. Interestingly, while complement was essential for most mAbs, immunization experiments revealed that A28 can also elicit complement-independent neutralizing antibodies. This dichotomy suggests that A28 harbors multiple neutralizing epitopes with diverse functional properties, highlighting the potential for inducing robust immunity through targeted antigen design. Another key finding is the synergy achieved by combining mAbs targeting A28 with anti-H3 or anti-A35 mAbs. Targeting different antigens enhanced neutralization breadth and potency, highlighting the value of multi-targeted therapeutic strategies.

Finally, our study demonstrates the feasibility of rational vaccine design based on A28, with vaccinated mice showing complete protection from a lethal VACV viral challenge. Vaccinations remain the most efficient method for preventing infections or complications from viral diseases. Historically, Poxviruses were the first pathogens to be controlled through immunization, culminating in the successful eradication of Smallpox in 1980^61^. While the non-replicating MVA vaccine strain mitigated viral spread^8^, its antigenic makeup differs from Mpox^45,62^, and the immune response rapidly declines over time^9-11^. The low immunogenicity of MVA, along with recent advances in vaccine technology, has facilitated the development of new antigen-based vaccines^63-66^. Recent studies have demonstrated that poxvirus mRNA-and subunit-based vaccines elicit robust neutralizing and protective antibody responses in both mouse and non-human primate (NHP) models^29,63-68^. These vaccine candidates primarily include antigens known to be targets of neutralizing antibodies and therefore typically exclude A28 as an immunogen. We show that immunization of mice with recombinant A28 elicits strong antigen-specific B cell responses and neutralizing antibodies, some of which functioned independently of complement. Interestingly, a common feature of the human anti-A28 mAbs identified in this study is that their activity was largely germline-encoded, and did not depend on somatic mutations introduced in their coding antibody sequences. This suggests that anti-A28 antibodies may be readily elicited without requiring extensive affinity maturation, which is an encouraging prospect for vaccine design.

### Limitations of the study

This study has several limitations. Due to the complexity of the Mpox virus and its abundance of surface proteins, we focused on a limited number of antigens (A35, H3, and A28) to assess the immune response, potentially overlooking other relevant targets that contribute to protection. A broader analysis including additional viral antigens could provide a more comprehensive understanding of the antibody response. Second, while we employed antigen-specific B cell sorting, some of the antigens used in our study, particularly H3 and A28, also serve as viral attachment proteins. This likely led to non-specific binding to B cells during sorting and may explain the small proportion of antigen-binding mAbs. As a result, we were unable to fully characterize the B cell repertoire responding to these antigens, potentially leading to an incomplete representation of the antibody diversity generated post-infection. Additionally, due to biosafety level restrictions, we primarily used VACV as a surrogate to study the mechanism of mAb neutralization rather than Mpox. While VACV shares many structural and functional similarities with Mpox, differences between the two viruses may limit the direct applicability of our findings to Mpox-specific immunity.

## SUPPLEMENTAL FIGURE LEGENDS

**Figure S1:**
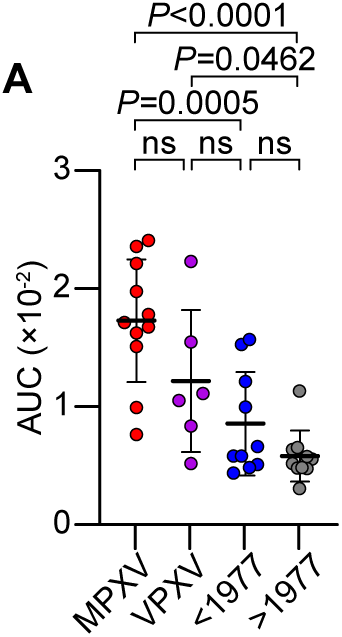
Antibody binding to A28 and VACV IHDJ. **(A)** Sera binding to Mpox A28 antigen as measured by ELISA. AUC values of 4 consecutive 4-fold dilutions, starting from 1:100. Mpox convalescent donors, MPXV, are in red (n=11), recently vaccinated donors, VPXV, are in purple (n=6), historic vaccinated donors born before 1977, <1977, are in blue (n=10) and naïve donors born after 1977, >1977, are in gray (n=10). Statistical analysis was performed using One-Way ANOVA with Tukey’s multiple comparison correction. ns= non significant.

**Figure S2:**
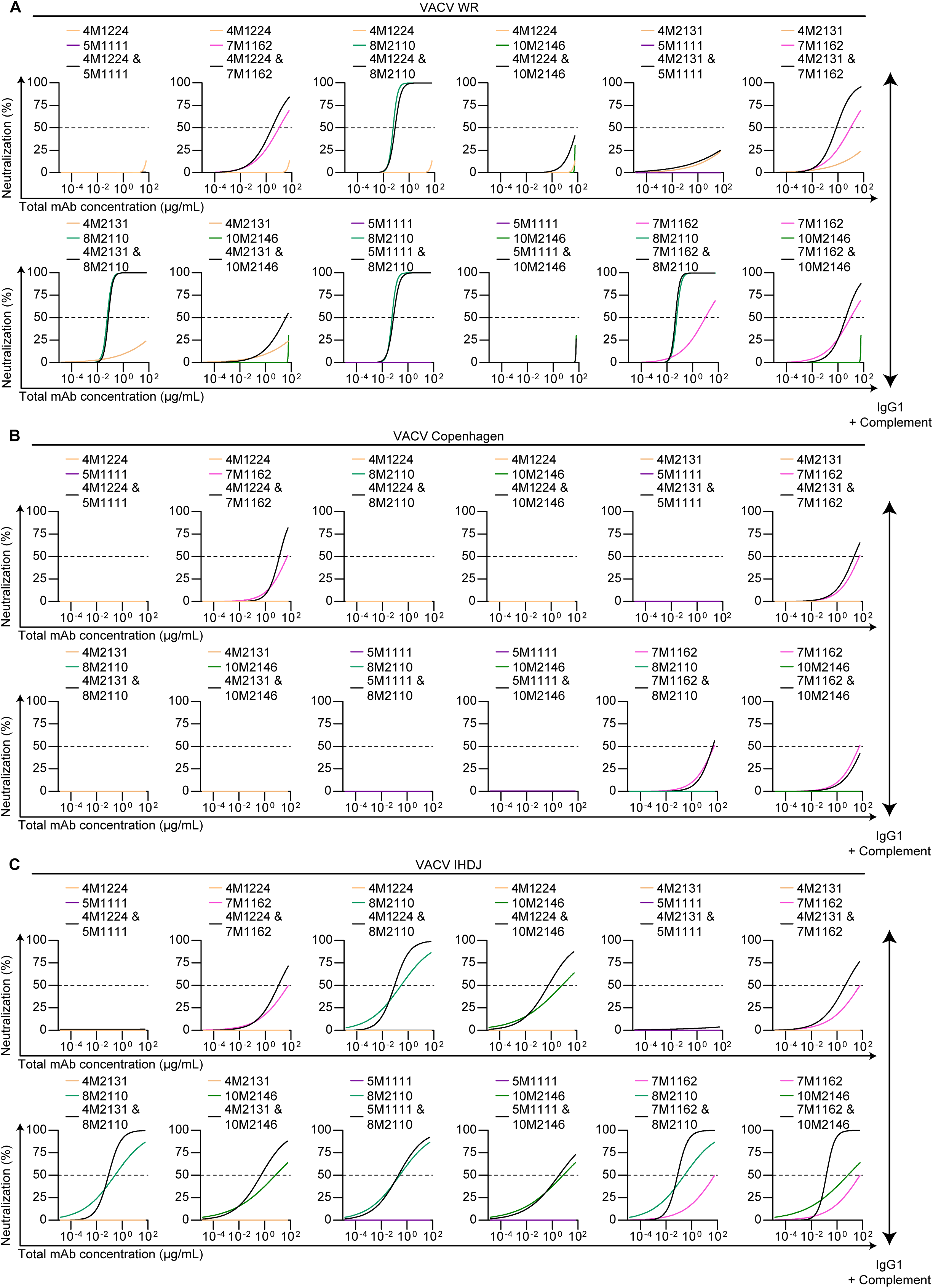
Increased potency of anti-Mpox mAb combinations. **(A)** Neutralization of VACV WR strain by anti-Mpox pairs in the presence of 1.5% NHS. Neutralization curves of 12 consecutive 4-fold dilutions, starting from 62.5 μg/mL. Each combination, as well as the individual mAbs, is depicted in a different panel. **(B)** Same as **A** but for VACV Copenhagen. **(C)** Same as **A** but for VACV IHDJ. Anti-Mpox mAbs are depicted in the following color scheme: anti-A35 mAbs in orange, anti-H3 mAbs are in magenta and purple and anti-A28 mAbs in shades of green. Combinations are in black. Neutralization curves were determined by fitting values using the Agonist vs. normalized response (Variable slopes) nonlinear regression.

**Figure S3:**
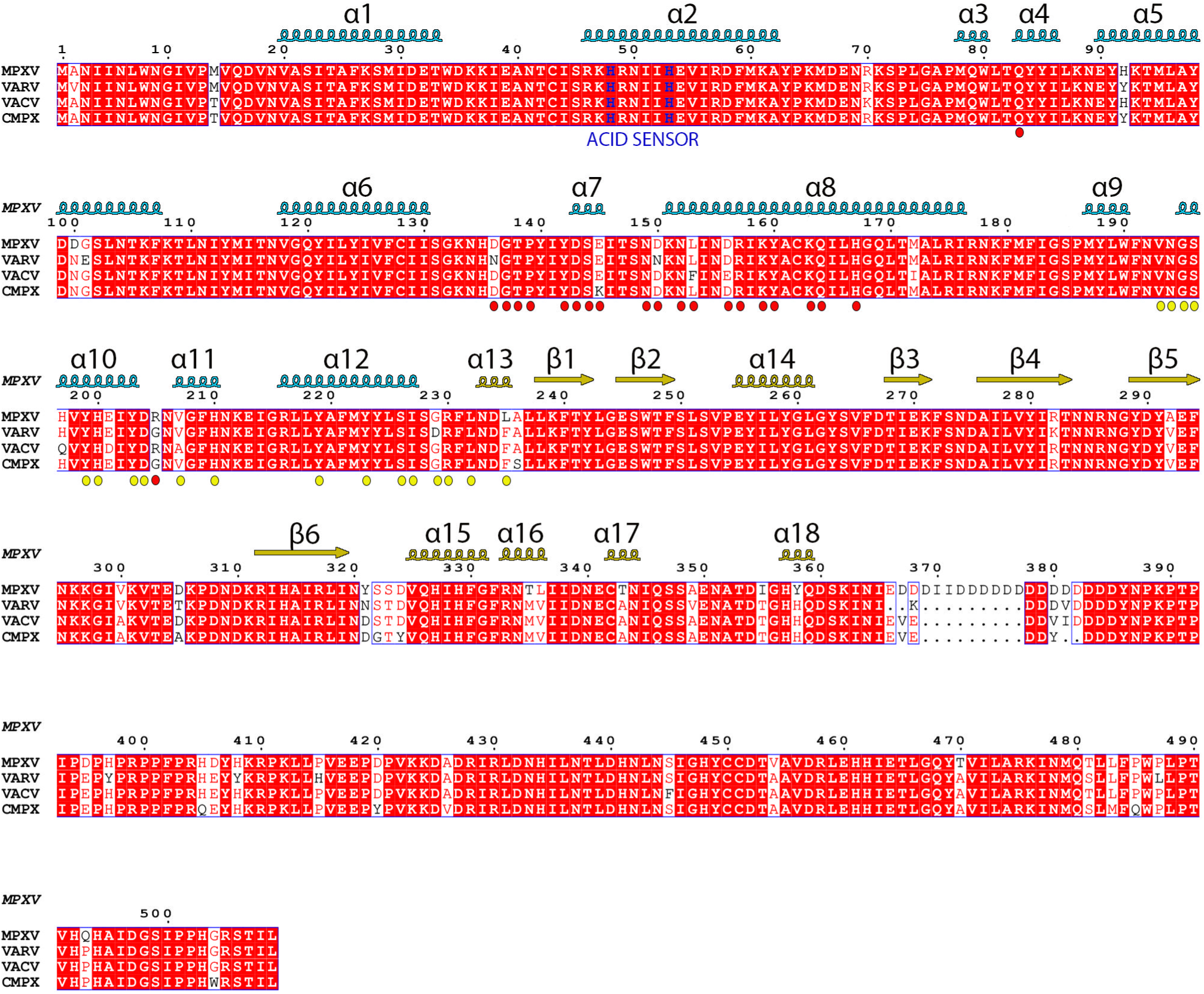
Multiple sequence alignment of A28 homologs. **(A)** A28 homologs (OPG153) from four representatives Orthopoxviruses—Mpox virus (MPXV; accession A0A7H0DNC4), Variola virus (VARV; Q89489), Vaccinia virus (VACV; P24758), and Camelpox virus (CMPX; Q8QQ29). Secondary structure elements are labeled above the sequences and colored by domain (NTD in blue, CTD in tan). Strictly conserved residues are highlighted with a red background. Epitope residues (BSA > 10 Å^2^) are marked by color-coded spheres below the sequences: red for 10M2146 and yellow for 8M2110.

**Figure S4:**
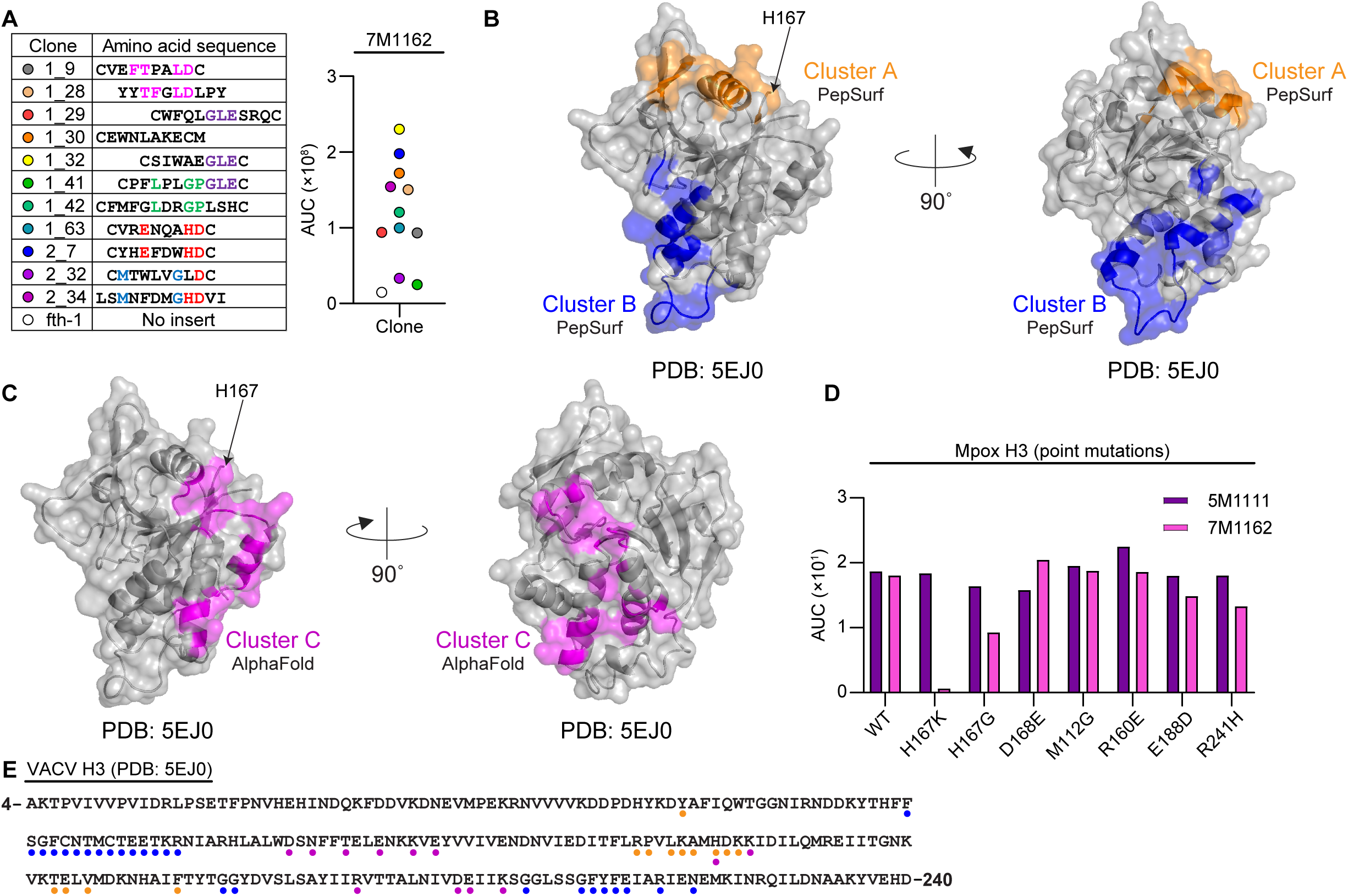
Prediction of the epitope of anti-H3 mAb 7M1162. **(A)** Left panel: peptide sequences isolated using Biopanning of a random phage display peptide library screened with the anti-H3 mAb 7M1162. Shared motifs are marked by matching colors. Right panel: The AUC values in ELISA of 8 consecutive 4-fold phage dilutions, starting from 4×10^8^ phages per well. fth-1 ( a phage without any recombinant peptide) serves as a negative control. **(B)** MAb 7M1162 predicted epitope on the surface of VACV H3 (PDB: 5EJ0) as predicted by Pepsurf software using isolated peptides. Predicted epitope clusters A and B are in orange and blue, respectively. **(C)** MAb 7M1162 predicted epitope on the surface of VACV H3 (PDB: 5EJ0) as predicted by AlphaFold3 software. Predicted epitope cluster C is in magenta. **(D)** Binding of 7M1162 (pink) and 5M1111 (purple) to Mpox H3 point mutations as measured by ELISA. AUC values of 6 consecutive 10-fold dilutions, starting from 10 μg/mL. **(E)** Sequence representation of predicted epitopes. Inferred epitope residues are marked by color-coded spheres below the sequences: orange for cluster A, blue for cluster B and magenta for cluster C.

**Figure S5:**
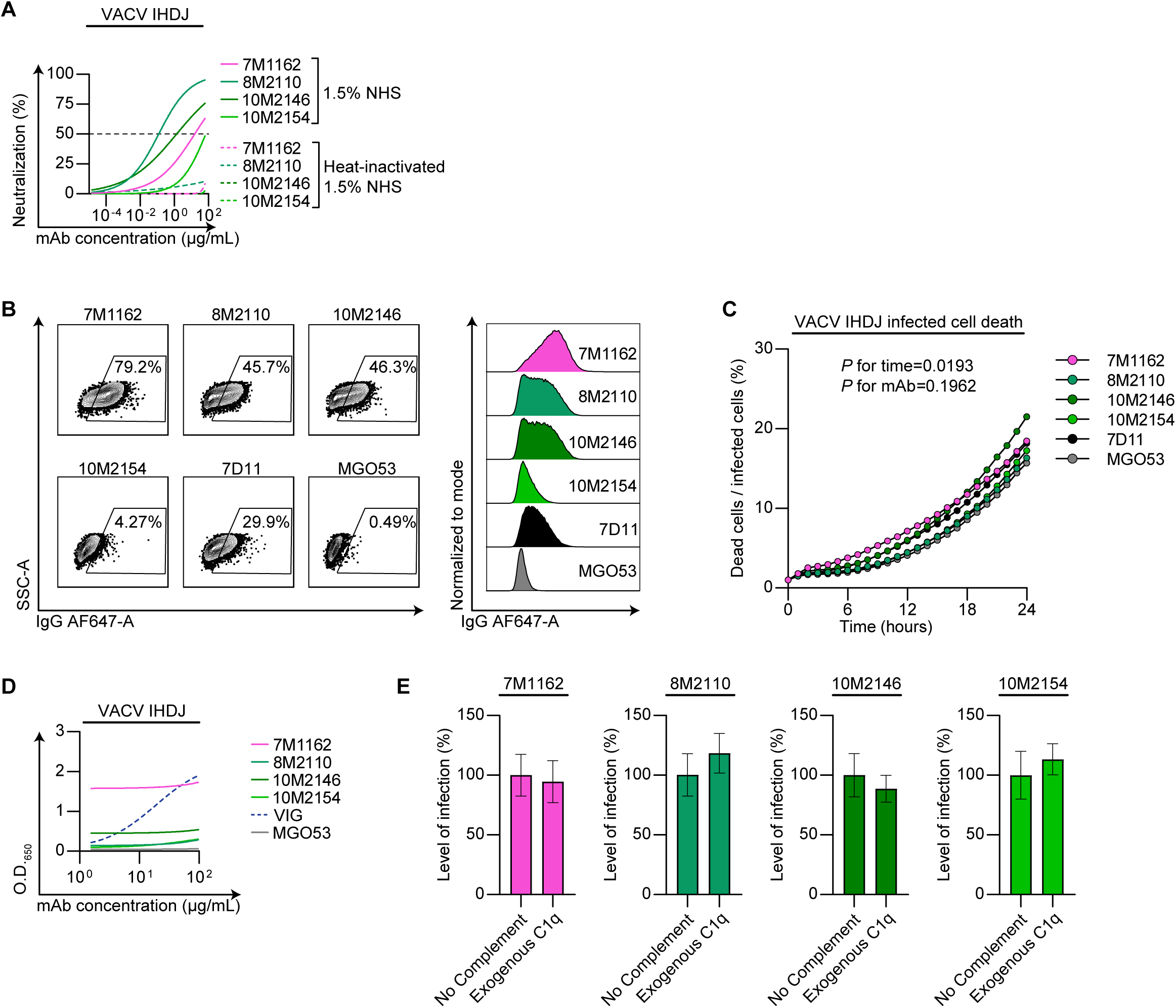
Anti-Mpox mAbs do not elicit CDC of VACV infected cells. **(A)** Neutralization of anti-Mpox mAbs against VACV IHDJ in the presence of 1.5% NHS versus in the presence of heat-inactivated NHS. Neutralization curves of 12 consecutive 4-fold dilutions, starting from 62.5 μg/mL. Dashed line represents 50% neutralization. **(B)** Binding of anti-Mpox mAbs [10 μg/mL] to VACV infected cells. Left panel: flow cytometry plots. Right panel: Median florescent intensity. **(C)** Induction of CDC on VACV infected cells. Baseline cell death was measured for each well at time point 0, following the addition of 10 μg/mL anti-Mpox mAbs in the presence of 1.5% NHS. Cell death was monitored over the next 24 hours using a cell death marker and live cell imaging. **(D)** Binding curves of anti-Mpox mAbs to VACV IHDJ as measured by ELISA of 4 consecutive 4-fold dilutions, starting from 100 μg/mL. **(E)** Level of VACV IHDJ infection in the presence of anti-Mpox mAbs in IC_50_ concentrations, with or without the addition of 1.2 μg/mL exogeneous C1q (compatible with levels found in 1.5% NHS). Anti-Mpox mAbs are depicted in the following color scheme: anti-H3 mAb 7M1162 is in magenta and anti-A28 mAbs are in shades of green. Vaccinia immune globulin (VIG) is in dashed blue, anti-M1 (7D11) as a positive control is in black and MGO53 as an isotype control is in gray. For the neutralization assays shown in panel **E**, the following mAb concentrations, corresponding to their IC_50_ values, were used: 15 μg/mL for 7M1162, 0.125 μg/mL for 8M2110, 1.25 μg/mL for 10M2146, and 62.5 μg/mL for 10M2154. Statistical analysis for **C** was performed using Two-Way repeated measures ANOVA with Dunnet’s multiple comparison correction to the MGO53 treated wells as control. Neutralization curves were determined by fitting values using Agonist vs. normalized response (Variable slopes) nonlinear regression for **A**. Binding curves were determined by fitting values using the Agonist vs response (three parameters) nonlinear regression for **B** Standard deviation of mean are shown in **E**.

**Figure S6:**
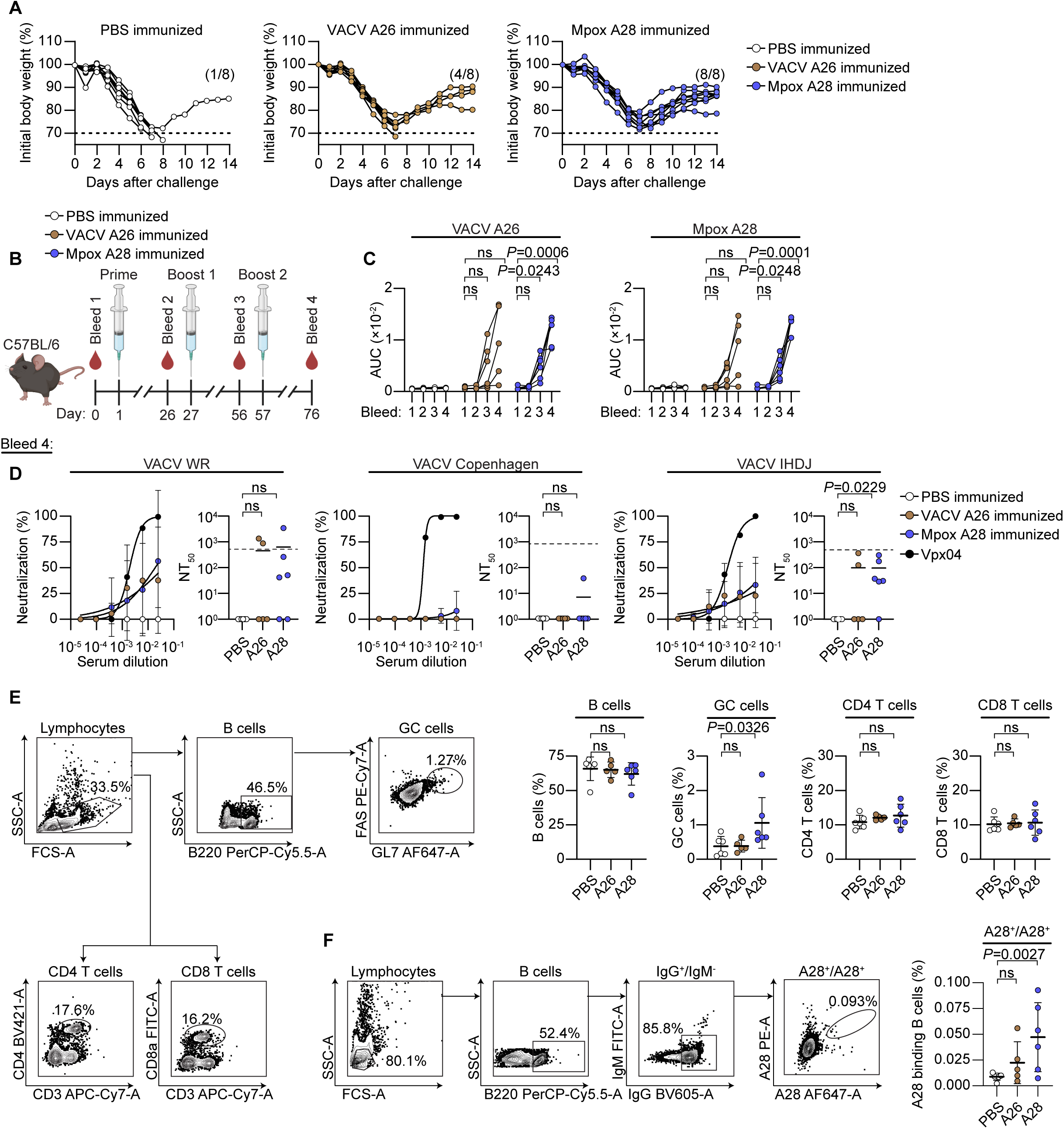
A28 vaccination elicits specific antibody and B cell responses. **(A)** Body weight changes for each immunized mouse from Figures 6D and **6F**. The number of mice that reached the experimental endpoint is indicated in parentheses. Left panel: PBS-immunized mice. Middle panel: VACV A26–immunized mice. Right panel: Mpox A28–immunized mice. **(B)** Schematic representation of mice immunization protocol. C57BL/c mice (n = 6-7 per group) were immunized I.P. with either PBS, Mpox A28, or VACV A26 antigens (10 μg per dose) formulated with alum. Immunizations were administered three times (prime + two boosts) at 3-4-week intervals. The day leading to each inoculation, the mice were bled, and their sera was isolated, and heat inactivated. All Mice were sacrificed 3 weeks after receiving the second boost with their blood and spleens harvested and analyzed. **(C)** Time course analysis of sera binding as measured by ELISA. AUC values of 4 consecutive 4-fold dilutions, starting from 1:100. Left panel: for VACV A26 antigen. Right panel: for Mpox A28 antigen. **(D)** Neutralization of sera at Bleed 4 (prime + 2 boosts) against VACV strains in the presence of 1.5% NHS. Neutralization curves of 6 consecutive 4-fold dilutions, starting from 1:40. NT_50_ values of neutralization curves are used for analysis. From left to right, panels represent neutralization of VACV WR, VACV Copenhagen and VACV IHDJ strains. **(E)** Flow cytometry analysis of splenocyte immune cells populations in vaccinated mice. Left panel: representative plots showing the gating strategy from an A28 vaccinated mouse. Right panels: percentage of total B cells, GC B cells, CD4 T cells and CD8 T cell population, as calculated for each vaccinated mouse. **(F)** Flow cytometry analysis of Mpox A28 binding B cells isolated from spleens of vaccinated mice. Left panel: representative plot and gating strategy from an A28 vaccinated mouse. Right panel: percentage of Mpox A28 binding B cells as calculated for each mouse. PBS immunized mice are in white (n=7 at initial starting point), VACV A26 immunized mice are in brown (n=7 at initial starting point) and Mpox A28 immunized mice are in blue (n=6 at initial starting point). Sera from a vaccinated individual (Vpx04), immunized three times with a VACV based vaccine is in black. For **B**, statistical analysis was performed by comparing mice sera AUC values to their baseline response from bleed 1 using Two-Way repeated measure ANOVA with Dunnett’s multiple comparison correction. For **C** and **D**, statistical analysis was performed using Kruskal-Wallis test with Dunn’s multiple comparison correction. For **E**, statistical analysis was performed using One-Way ANOVA with Dunnet’s multiple comparison correction. Images were created using BioRender. ns= non significant.

## RESOURCE AVAILABILITY

### Lead contact

Further information and requests for resources and reagents should be directed to and will be fulfilled by the lead contact, Natalia T Freund.

### Materials availability

Monkeypox virus antigens generated in this study will be available upon request from the lead contact with a completed Materials and Transfer Agreement.

### Data and code availability

- All data reported in this paper will be shared by the lead contact upon request.
- This paper does not report original code.
- Any additional information required to reanalyze the data reported in this paper is available from the lead contact upon request

## EXPERIMENTAL MODEL AND SUBJECT DETAILS

### Ethics statement

Peripheral whole blood samples were collected at 1–2 months and 9–10 months post-Mpox infection or primary vaccination. All Mpox convalescent and vaccinated donors provided written informed consent before sample collection at each time point. The Tel Aviv University Institutional Review Board (IRB) approved all studies involving patient enrollment, sample collection, and clinical follow-up (protocol number 0005243–1; Helsinki approval number 0384-22-TLV). In vivo studies complied with Tel Aviv University-Institutional Animal Care and Use Committee (permit number TAU - MD - IL - 2407 - 149 – 4, TAU - MD - IL - 2411 - 181 – 5 and TAU - MD - IL - 2502 - 104 - 5).

### Cell lines and viruses

Expi293F cells (Thermo Fisher Scientific) were grown in Expi293 Expression Medium (Thermo Fisher Scientific) at a constant shaking speed of 110 RPM and maintained at 37°C with 8% CO₂. HeLa (human female cervical cancer, ATCC CCL-2), Vero (African green monkey kidney epithelial cells, ATCC CCL-81), and U2OS (human female osteosarcoma cells, ATCC HTB-96) cells were cultured in DMEM (Sigma-Aldrich) supplemented with 10% heat-inactivated (HI) FBS (Hyclone), 100 U/mL penicillin, 100 U/mL streptomycin, and 2 mM L-glutamine (Thermo Fisher Scientific). Cells were maintained at 37°C with 5% CO₂.

The VACV IHDJ strain used for ELISA was kindly provided by Prof. Ehud Katz and was inactivated using β-propiolactone (βpL) by the Israel Institute of Biological Research (IIBR) prior to use under BSL-1 conditions. VACV-vFIRE-WR, VACV-vFIRE-Copenhagen, and VACV-vFIRE-IHDJ were kindly provided by Prof. Bernard Moss. The Mpox strain (MPXV/2022/FR/CMIP) was isolated and propagated as previously described^42,69^.

VACV-vFIRE-WR, VACV-vFIRE-Copenhagen, and VACV-vFIRE-IHDJ were propagated on a confluent monolayer of HeLa cells in MEM (Gibco) supplemented with 2% HI FBS, 100 U/mL penicillin, 100 U/mL streptomycin, 2 mM L-glutamine, and 1% non-essential amino acids (NEAAs) (Sartorius). After 2 hours, the cells were overlaid with MEM supplemented with 8% HI FBS, 100 U/mL penicillin, 100 U/mL streptomycin, 1% NEAAs, and 2% sodium bicarbonate solution (Merck). After 72 hours of virus propagation, infected cells were scraped from the tissue culture dish and pelleted by centrifugation at 1,800 × g for 5 minutes. Viruses were harvested by three freeze-thaw cycles, and cell debris was removed by centrifugation at 1,800 × g for 5 minutes. Viruses were titrated on a confluent monolayer of Vero cells. Briefly, Vero cells were infected with serially diluted virus. After 1 hour of infection, the cells were overlaid with DMEM containing methylcellulose (Sigma-Aldrich) and incubated for 72 hours, followed by plaque visualization using Coomassie staining (Bio-Rad). For experiments requiring purified MV particles, infected cell lysates were centrifuged and overlaid onto a 36% sucrose cushion, followed by ultracentrifugation at 33,000 × g for 80 minutes. The resulting pellet was then resuspended in 10 mM Tris-HCl (pH 9)

All experiments with infectious VACV strains were performed under BSL-2 or ABSL-2 and conditions. All experiments with infectious Mpox were performed under strict BSL-3 conditions.

### Mice

Female C57BL/6JOlaHsd and BALB/cOlaHsd mice were purchased at 7 weeks of age from Harlan Laboratories. Experimentation complied with Tel Aviv University-Institutional Animal Care and Use Committee (permit number TAU - MD - IL - 2407 - 149 – 4, TAU - MD - IL - 2411 - 181 – 5 and TAU - MD - IL - 2502 - 104 - 5).

## METHOD DETAILS

### Sample collection and processing

Peripheral whole blood samples were collected at 1–2 months (time point 1, V1) and 9–10 months (time point 2, V2) following PCR-confirmed Mpox infection or primary vaccination with the JYNNEOS vaccine. Serum was isolated from 5 mL of each sample, while the remaining whole blood was used for peripheral blood mononuclear cell (PBMC) isolation using a Ficoll gradient (Cytiva), as previously described^33^. For serum isolation, whole blood was allowed to clot and then centrifuged at 2,000 × g for 15 minutes, followed by heat inactivation at 56°C for 20 minutes.

### Expression of Mpox antigens and VACV homologs

Expression of recombinant Mpox antigens A35 and H3 was previously described^17^. Mpox (GenBank: MN648051)^70^ and VACV IHDW strain (GenBank: KJ125439)^71^ sequences were used as templates for the production of Mpox antigen A28 (OPG 153) and the VACV homologs A33 (OPG 161), H3 (OPG 108), and A26 (OPG 153). The amino acid regions expressed for each antigen were as follows: Mpox A28 (2–361), VACV A33 (58–185), VACV H3 (2–278), and VACV A26 (2–361). Construct design was similar to that of Mpox antigens A35 and H3^17^. Briefly, antigen amino acid sequences were codon-optimized for mammalian expression and modified to include a growth factor receptor signal peptide at the N-terminus. An 8×Histidine (His-tag) sequence followed by a biotinylation encoding sequence (Avi-tag) was added at the C-terminus. Antigen-encoding DNA sequences were synthesized by an external vendor, cloned into the pcDNA3.1 (-) vector, and transiently transfected into Expi293F cells following the manufacturer’s protocol. Seven days post-transfection, cell supernatants were incubated with nickel beads (Cytiva) overnight at 4°C. The beads were then loaded onto chromatography columns (Bio-Rad), washed, and the antigens were eluted using increasing concentrations of imidazole (Thermo Scientific), followed by dialysis against PBS overnight. For experiments requiring biotinylated antigens, buffer exchange was performed with 10 mM Tris-HCl, and biotinylation was carried out using the BirA kit (Avidity), following the manufacturer’s protocol. Biotinylated antigens were then buffer-exchanged back to PBS.

Mpox H3 point mutations were generated by PCR amplification of the Mpox H3 plasmid with corresponding nucleotide substitutions and cloned as previously described^17^. H3 mutations were verified by Sanger sequencing, and the mutant H3 proteins were produced in Expi293F cells as described above.

### Western blot analysis of purified antigens

Validation of proper expression of Mpox and VACV antigens was performed using SDS-PAGE and Western blotting. Briefly, 0.5 μg of each antigen, containing 25 mM DTT (Sigma-Aldrich), along with an unstained protein standard (Bio-Rad), were loaded onto an SDS-PAGE and transferred to a nitrocellulose membrane (Bio-Rad). The membrane was blocked for 2 hours at room temperature (RT) in blocking buffer containing PBS, 3% BSA (ENCO), 0.05% Tween-20 (Sigma-Aldrich), and 20 mM EDTA (Bio-Labs). Anti-Avi-Tag mouse antibody (Avidity), diluted 1:5000 in blocking buffer, was incubated with the membrane overnight at 4°C. After 5 washes with washing buffer containing PBS and 0.05% Tween-20, the membrane was incubated for 45 minutes at RT with a secondary anti-mouse HRP-conjugated antibody (Jackson Laboratory) and StrepTactin-HRP (Bio-Rad), both diluted 1:5000 in blocking buffer. Following 5 additional washes, ECL substrate was added, and Western blot images were acquired in Chemidoc imaging system (Bio-Rad).

### Antigen-binding single B cell sorting and sequencing

PBMCs from Mpox convalescent donors were thawed in RPMI 1640 (Sigma-Aldrich) medium warmed to 37°C, followed by centrifugation and resuspension in MACS buffer containing PBS, 0.5% BSA, and 2 mM EDTA. B cells were then enriched using anti-CD19 (Miltenyi Biotec) magnetic beads according to the manufacturer’s protocol, followed by centrifugation and resuspension in FACS buffer containing PBS, 1% FBS, and 2 mM EDTA for subsequent staining. The B cell-enriched samples were stained with Labeling Check Reagent-VioBlue (Miltenyi Biotec), IgG-FITC (BioLegend), and either biotinylated Mpox antigens A35, H3, or A28, labeled with both Streptavidin-PE (BioLegend) and Streptavidin-Alexa Fluor 647 (BioLegend), for 30 minutes at 4°C. After washing and resuspending the cells in FACS buffer, CD19^+^IgG^+^antigen^+^ cells were single-cell sorted using the FACSAriaIII sorter into 96-well PCR plates containing 4 μL of lysis buffer (PBS×0.5, 12 units of RNasin Ribonuclease inhibitor (IMBH), and 10 mM DTT (Invitrogen), and immediately frozen on dry ice. Plates were thawed on ice for subsequent cDNA synthesis and PCR amplification as previously described^33,72^. Briefly, lysed cells were prepared for reverse transcription by adding random hexamer primers (Invitrogen), IGEPAL (Sigma-Aldrich), and RNasin Ribonuclease Inhibitor to the sorted wells and incubating at 68°C for 1 minute. cDNA was then synthesized using a reverse transcription PCR reaction containing SuperScript III reverse transcriptase (Invitrogen), dNTPs (Thermo Scientific), DTT, and RNasin Ribonuclease Inhibitor. The reaction was carried out under the following conditions: 42°C for 10 minutes, 25°C for 10 minutes, 50°C for 60 minutes, and 94°C for 5 minutes. Assembled cDNA was used as a template for subsequent Ig_H_/Ig_L_ amplification by two-step nested PCR. The first round of PCR was performed using Phusion High-Fidelity DNA Polymerase (NEB), dNTPs, and the corresponding immunoglobulin Gamma, Kappa, or Lambda primer mix^33,72^. The reaction conditions were as follows: initial denaturation at 98°C for 30 seconds, followed by 50 cycles of denaturation at 98°C for 10 seconds, annealing at 52°C (Gamma), 50°C (Kappa), or 58°C (Lambda) for 15 seconds, extension at 72°C for 15 seconds, and final extension at 72°C for 5 minutes. The second PCR was performed under similar conditions, but the number of cycles was reduced to 40, and the annealing temperatures were adjusted to 56°C (Gamma), 50°C (Kappa), and 60°C (Lambda). To verify DNA amplification, the second-round PCR products were visualized on a 2% agarose gel and sent for Sanger sequencing.

The retrieved Ig_H_/Ig_L_ sequences were analyzed and annotated using IgBLAST and the IMGT database for sequence quality, viability, Gamma chain subtype (Ig_H_ only), V_H_D_H_J_H_ or V_L_J_L_ gene usage, mutational load, and CDR3 length. Cells displaying identical V and J gene usage and ≥70% homology in both Ig_H_/Ig_L_ sequences were classified as clonally expanded B cells. Germline sequences for each mAb were determined using the IMGT database. For Ig_H_ CDR3, the germline sequence was set as the retrieved Ig_H_ CDR3 sequence itself. In cases where clonally expanded B cells were identified, both CDR3 Ig_H_/Ig_L_ sequences were determined by assessing the most likely common features.

### Anti-Mpox antibody cloning and production

Viable Ig_H_/Ig_L_ sequences recovered from previous steps were selected for production as IgG1 mAbs. Briefly, first-round PCR products were used as templates for an additional round of PCR amplification with specific 5′ and 3′ primers encoding restriction sites for subsequent cloning. This additional PCR was performed under conditions similar to the previously described second-round PCR, with an annealing temperature of 56°C for all reactions. PCR products were visualized on a 2% agarose gel, purified, digested with the appropriate restriction enzymes (NEB), and ligated into the relevant immunoglobulin expression vectors: IgG1 (Gamma), IgK (Kappa), and IgL (Lambda). The corresponding heavy (Gamma) and light (Kappa or Lambda) chain plasmids were transiently co-transfected at a 1:3 ratio into Expi293F cells following the manufacturer’s protocol. Seven days post-transfection, the supernatant was incubated with protein A beads (Cytiva) for 2 hours at room temperature (RT). Beads were then loaded onto chromatography columns, washed, and mAbs were eluted using a low-pH solution before being dialyzed against PBS overnight. For experiments requiring biotinylated mAbs, biotinylation was performed using EZ-Link Sulfo-NHS-LC-Biotin (Thermo-Scientific) according to the manufacturer’s protocol, followed by an additional overnight dialysis against PBS. To produce germline-reverted mAbs, codon-optimized sequences of the germline-reverted mAbs (as described above) were synthesized by an external vendor and cloned into the relevant immunoglobulin expression vector. Plasmids were transiently co-transfected into Expi293F cells, and germline-reverted mAbs were isolated as previously described.

Anti-VACV-L1 mouse recombinant mAb (7D11)^73^ was used as a positive control for subsequent neutralization assays, as previously described^25,65^. Briefly, the Ig_H_/Ig_L_ sequences of 7D11 were obtained from the Protein Data Bank (PDB: 2I9L)^74^. The corresponding amino acid sequences were codon-optimized, synthesized by an external vendor, and cloned into a human IgG1 and IgK immunoglobulin expression vectors. Cloned plasmids were transiently co-transfected into Expi293F cells, and the humanized 7D11 mAb was purified as previously described. The 7D11 mAb was first tested against the Mpox M1 antigen^17^ in ELISA before being used as a positive control in the subsequent neutralization assays (data not shown).

### Enzyme-linked immunosorbent assay

VACV IHDJ ELISA: βpL-inactivated VACV IHDJ strain was diluted in carbonate-bicarbonate solution to a concentration of 10L PFU/mL and used to coat high-binding ELISA plates (Corning) overnight at 4°C. Plates were then blocked for 2 hours at RT with blocking buffer comprising PBS, 3% BSA, 0.05% Tween-20, and 20 mM EDTA, followed by a single wash with washing buffer comprising PBS and 0.05% Tween-20. Anti-Mpox mAbs were serially diluted 4-fold in blocking buffer, beginning at 100 μg/mL, for a total of four dilutions before being added to the plates for 1 hour at RT. After three washes with washing buffer, plates were incubated for 45 minutes at RT with an anti-human IgG HRP-conjugated antibody (Jackson Laboratory) diluted 1:5000 in blocking buffer. Following five additional washes, TMB/E (Abcam) was added, and optical density (O.D.) was measured at 650 nm after 10 minutes.

Mpox and VACV antigen ELISA: 2.5 μg/mL of antigen was used to coat high-binding ELISA plates overnight at 4°C. Plates were then blocked for 2 hours at RT with a blocking buffer, followed by a single wash with washing buffer. For the Mpox convalescent and vaccinated donor ELISA, HI serum samples were serially diluted 4-fold in blocking buffer, starting at 1:100, for a total of four dilutions. Alternatively, for anti-Mpox mAb ELISA, mAbs were diluted in blocking buffer across 12 consecutive 4-fold dilutions, starting at 10 μg/mL. Serum or mAb samples were added to the ELISA plates and incubated for 1 hour at RT. After three washing cycles with washing buffer, plates were incubated with an anti-human IgG HRP-conjugated antibody (diluted 1:5000 in blocking buffer) for 45 minutes at RT. After five additional washes, TMB/E was added, and the O.D. was measured at 650 nm after 10 minutes.

Immunized mice ELISA: 2.5 μg/mL of antigen was used to coat high-binding ELISA plates overnight at 4°C. Plates were then blocked for 2 hours at RT with a blocking buffer, followed by a single wash with washing buffer. HI vaccinated mouse serum was serially diluted 4-fold in blocking buffer, starting at 1:100 and added to the ELISA plates and incubated for 1 hour at RT. After three washing cycles with washing buffer, plates were incubated with an anti-mouse IgG HRP-conjugated antibody (Jackson Laboratory) (diluted 1:5000 in blocking buffer) for 45 minutes at RT. After five additional washes, TMB/E was added, and the O.D. was measured at 650 nm after 10 minutes.

Anti-Mpox mAbs competition ELISA: 1 μg/mL of Mpox A35 and H3, and 2.5 μg/mL of Mpox A28, were used to coat high-binding ELISA plates overnight at 4°C. The plates were then blocked for 2 hours at RT in blocking buffer, followed by a single wash with washing buffer. The anti-Mpox mAbs were diluted in blocking buffer across 8 consecutive 4-fold dilutions, starting at 64 μg/mL, and added to the ELISA plates for 30 minutes at RT. Next, 0.1 μg/mL of biotinylated mAbs (4M1130, 4M1133, or 4M1154 for A35-coated plates, 5M1111 or 7M1162 for H3-coated plates, and 8M2110 or 10M2146 for A28-coated plates) were added to each well and incubated for an additional 15 minutes. The plates were then washed three times with washing buffer and incubated with Streptavidin-HRP (Jackson Laboratory), diluted 1:5000 in blocking buffer, for 45 minutes at RT. After 5 additional washes, TMB/E was added, and the O.D. was measured at 650 nm after 10 minutes.

Affinity-selected phage ELISA: 2 μg/mL of mAbs 4M1130, 4M1224, and 7M1162 were used to coat high-binding ELISA plates overnight at 4°C. The plates were then blocked for 2 hours at RT in a blocking solution comprising TBS, 5% skim milk (Difco), and 0.05% Tween20, followed by a single wash with washing solution comprising TBS and 0.05% Tween20. Corresponding affinity-selected phages for each mAb were diluted across 8 consecutive 4-fold dilutions in blocking solution, starting from 4×10^8^ phages per well, and added to the ELISA plates for 1 hour at RT. The plates were then washed three times with washing solution and incubated with rabbit anti-M13 antibody diluted 1:5000 in blocking solution for 1 hour at RT. After 3 additional washes, the plates were incubated with anti-rabbit IgG HRP-conjugated antibody (Merck) diluted 1:5000 in blocking solution for 45 minutes at RT. Following 5 more washes, TMB/E was added, and the O.D. was measured at 650 nm after 10 minutes.

### Virus neutralization assays

VACV strains: WR (VACV-vFIRE-WR), Copenhagen (VACV-vFIRE-Copenhagen), and IHDJ (VACV-vFIRE-IHDJ), as well as Mpox clade IIb (MPXV/2022/FR/CMIP), were used for antibody neutralization assays. All neutralization assays were performed on U20S cells, which were seeded at 2×10^4^ cells per well in a 96-well plate 24 hours before virus infection.

VACV WR, Copenhagen, and IHDJ strains neutralization assay: Anti-Mpox mAbs or HI mouse sera were diluted in 6 consecutive 4-fold dilutions in PBS, starting from 250 μg/mL and 1:25, respectively. For experiments requiring IC_50_ determination, mAbs were diluted in 12 consecutive 4-fold dilutions. Antibodies were mixed at a 1:1 ratio with cell medium containing 4×10³ PFU/mL of either VACV WR, Copenhagen, or IHDJ in the presence or absence of 6% Normal Human Serum (NHS) (Sigma-Aldrich) or HI NHS. After a 2-hour incubation at 37°C, the virus inoculum was added to the U2OS cell monolayer at a 1:1 ratio. The virus-infected wells had a final virus concentration of ∼100 PFU/well (MOI of 0.005), 1.5% NHS, 4-fold dilutions of mAbs starting from 62.5 μg/mL, or 4-fold dilutions of mouse sera starting from 1:100. For experiments in which anti-Mpox mAbs were tested in combinations, mAbs were mixed at a 1:1 ratio between themselves, reaching a final IgG concentration of 62.5 μg/mL (31.25 μg/mL of each mAb) and subsequently serially diluted in PBS if needed. Following incubation for 48 hours at 37°C and 5% CO2, plates were sealed with breathable sealing tape (Thermo Scientific), and images were acquired using the IncuCyte live-cell analysis system. Virus infection levels were quantified by analyzing the GFP area in each well.

Mpox neutralization assay: The neutralization assays using Mpox clade IIb were performed as previously described^42,75^. Anti-Mpox mAbs were diluted 6 consecutive 4-fold dilutions in PBS, starting from 250 μg/mL. For experiments requiring IC_50_ determination, mAbs were diluted 8 consecutive 4-fold dilutions starting from 200 μg/mL. MAbs were mixed at a 1:1 ratio with cell medium containing Mpox in the presence or absence of 40% Guinea pig complement (GPC-Rockland). After 2 hours of incubation at 37°C, virus inoculum was added onto the U20S cells monolayer at a 1:1 ratio. The virus-infected wells had a final concentration of 10% GPC, and 6 or 8 consecutive 4-fold dilutions of mAbs, starting from 62.5 μg/mL or 50 μg/mL, respectively. For experiments in which anti-Mpox mAbs were tested in combinations, mAbs were mixed at a 1:1 ratio between themselves, reaching a final IgG concentration of 62.5 μg/mL (31.25 μg/mL of each mAb). Following incubation for 48 hours at 37°C and 5% CO2, cells were fixed for 30 minutes with 4% PFA (Invitrogen), washed, and stained with rabbit polyclonal anti-VACV antibodies (Invitrogen). Following this, plates were stained with an anti-rabbit antibody coupled to Alexa Fluor 488 (Invitrogen) and washed with Hoechst (Invitrogen) diluted in PBS to visualize nuclei. Images were acquired with an Opera Phenix high-content confocal microscope, and infection levels were quantified by analyzing the fluorescence area in each well as previously described^42^.

Neutralization kinetics assay: VACV IHDJ neutralizing mAbs were used in their IC_50_ values: 15 μg/mL for 7M1162, 0.125 μg/mL for 8M2110, 1.25 μg/mL for 10M2146, and 62.5 μg/mL for 10M2154 in the presence of 1.5% NHS (final concentrations in the virus-infected wells). All wells were infected with 100 PFU/mL of VACV IHDJ simultaneously. The addition of diluted mAb and NHS were performed in 10 minutes intervals, starting 2 hours pre-infection and lasting until 2 hours post-infection. For pre-infection wells, mAbs and NHS were mixed in wells containing cell medium and ∼100 PFU/well of VACV IHDJ prior to its infection of the U2OS cells. For post-infection wells, mAbs and NHS were added directly to the previously infected cells. Analysis of neutralization was performed similarly to VACV neutralization assays described above.

Complement inhibitors and depleted sera neutralization assays: VACV IHDJ neutralizing mAbs were used at their IC_50_ values: 15 μg/mL for 7M1162, 0.125 μg/mL for 8M2110, 1.25 μg/mL for 10M2146, and 62.5 μg/mL for 10M2154. For experiments requiring complement depleted sera, mAbs were incubated in the presence of 1.5% NHS, C1q-, C3-depleted sera (Complement Technology, Inc., final concentrations in the virus-infected wells). For experiments involving selective complement inhibitors, mAbs and NHS mixtures were incubated with either 6 consecutive 4-fold dilutions, starting from 10 μM of a selective complement C5 inhibitor, C5-IN-1 (Compound 7-MedChemExpress), or 8 consecutive 5-fold dilutions, starting from 10 μM of a selective complement C3 inhibitor, CP40 (AMY-101-MedChemExpress). Analysis of neutralization was performed similarly to VACV neutralization assays described above.

Exogenous C1q neutralization assays: VACV IHDJ neutralizing mAbs were used at their IC_50_ values: 15 μg/mL for 7M1162, 0.125 μg/mL for 8M2110, 1.25 μg/mL for 10M2146, and 62.5 μg/mL for 10M2154 in the presence or absence of 1.2 μg/mL of human purified C1q, representing the amount found in 1.5% NHS (final concentrations in the virus-infected wells). Analysis of neutralization was performed similarly to VACV neutralization assays described above. The level of infection in samples containing C1q was normalized to those without.

In all virus neutralization assays, except where noted otherwise, the percentage of neutralization was calculated based on the reduction of fluorescence area (GFP for VACV or Alexa Fluor 488 for Mpox). Due to infection variability between wells, to ensure antibody-treated wells show genuine fluorescence area reduction, all non-antibody-treated wells were analyzed for their fluorescence area values. Baseline infection for each experiment was set as the non-antibody-treated well displaying the least fluorescence area (baseline infection). The percentage of neutralization for each well was calculated using the following formula: 100 - (fluorescence area value / baseline infection fluorescence area) × 100. Neutralization rates calculated as negative values were set to zero.

### VACV infected cells binding assay

Confluent monolayers of Vero cells were infected with VACV IHDJ (MOI of 1) for 14 hours until GFP expression became evident. Infected cells were then detached from the tissue culture dish using Trypsin/EDTA (Thermo Scientific), centrifuged at 400 × g for 5 minutes, and resuspended in FACS buffer containing 10 μg/mL of the anti-Mpox mAbs. The suspension was incubated for 30 minutes at 4°C. Following incubation, cells were washed, centrifuged again at 400 × g for 5 minutes, and resuspended in FACS buffer containing anti-IgG-Alexa Fluor 647 (BioLegend) for an additional 30 minutes at 4°C. Labeled cells were washed one more time and fixed with 4% PFA for 20 minutes at RT. After a final wash, the labeled cells were analyzed using a Flow Cytometer.

### VACV infected cells CDC assay

U20S cells were seeded at 2×10^5^ cells per well in a 6-well plate and incubated at 37°C and 5% CO2 for 24 hours until reaching ∼50% confluency. The cells were then infected with VACV IHDJ (MOI of 0.1) for an additional 24 hours, during which GFP expression became evident. To remove free viral particles, infected cells were washed twice with fresh media containing 1:1000 Propidium iodide (PI-BioLegend). Plates were sealed with breathable sealing tape, and the number of dead/total cells was analyzed using the IncuCyte live-cell analysis system at time point 0. After image acquisition, 10 μg/mL of anti-Mpox mAbs in the presence of 1.5% NHS were added to the wells containing infected cells. Plates were re-sealed, and further analysis of dead/total cells was conducted using the IncuCyte system. Images were acquired over the next 24 hours at 1-hour intervals.

### VACV C1q deposition

VACV IHDJ virions (∼5×10^6^ PFU) were incubated with 62.5 μg/mL of anti-Mpox mAbs and 1.5% NHS for 2 hours at 37°C. Following incubation, the virions were stained with anti-C1q-Alexa Fluor 647 (Santa Cruz) for 1 hour. PFA was then added to the virions to reach a final concentration of 4%, followed by analysis of the GFP-expressing virions in the Flow Cytometer.

### Immunogold labeling and transmission electron microscopy

Carbon/Formvar-coated copper grids (Bar Naor) were placed on a 40 µL drop of purified VACV IHDJ MV (5 × 10L PFU/mL) and incubated at room temperature for 15 minutes to facilitate virus adsorption. The grids were washed once with PBS and subsequently incubated in blocking solution (3% BSA in PBS) for 30 minutes. Following blocking, grids were incubated overnight at 4°C with 10 μg/mL of anti-Mpox mAbs in the presence of 1.5% NHS in blocking solution. The next day, grids were transferred to 37°C to allow complement activation, followed by two PBS washes and a 10-minute incubation in blocking solution. For immunogold labeling, grids were incubated for 3 hours at room temperature with either anti-C1q (Santa Cruz) or anti-C3 (BioLegend) in blocking solution, followed by additional washing and blocking steps. Secondary labeling was performed using anti-human IgG gold nanoparticles (10 nm) (Abcam) to visualize anti-Mpox mAbs and anti-mouse gold nanoparticles (20 nm) (Abcam) for C1q or C3 detection. Unbound antibodies were removed by two consecutive PBS washes (10 minutes each). Samples were fixed with 1% glutaraldehyde (Sigma-Aldrich) in PBS, rinsed with ultrapure water, and negatively stained with Uranyless (Electron Microscopy Sciences) according to the manufacturer’s instructions. The grids were air-dried and imaged using a JEM-1400 transmission electron microscope (JEOL Ltd, Tokyo, Japan) operating at 80 kV.

### Mice immunization

Passive immunization: Female BALB/c mice (8 weeks old) received 200 μg of either monoclonal antibody 8M2110 (anti-A28) or the isotype control MGO53, administered intraperitoneally in PBS. The following day mice were challenged.

Active immunization: For VACV challenge experiments, female BALB/c mice (8 weeks old) were immunized intraperitoneally with 10 μg of either Mpox A28, VACV A26, or PBS, formulated with 1.5Lμg of QS-21 adjuvant (InvivoGen) per mouse. Immunizations were administered three times (prime plus two boosts) at 3-week intervals. Two weeks after each injection, blood was collected and sera was heat inactivated. Three weeks after the final boost, mice were challenged. For immune cell profiling, female C57BL/6J mice (8 weeks old) were immunized intraperitoneally with the same antigen formulations (10 μg per dose), adjuvanted with Alum (Thermo Scientific) in a 1:1 volume ratio (final volume 200 μL). Immunizations were administered on days 1, 26, and 56. Mice were bled one day before each dose, and sera were collected and heat inactivated. On day 76, mice were sacrificed, and blood and organs were harvested for downstream analyses.

### VACV challenge experiments

BALB/c mice, either from the passive or active immunization experiments were housed under ABSL-2 conditions. Mice were anesthetized using Ketamine/Xylazine and inoculated I.N. with a lethal dose of VACV WR (2 × 10^5^ PFU). Weight and survival were monitored daily for 14 days. Mice that lost more than 30% of their initial weight (or more than 25% and exhibited a core body temperature below 34°C) were considered to have reached the no recovery threshold and were subsequently sacrificed. For viral load analysis, mice were inoculated with VACV WR as described above and sacrificed on day 5 post-infection.

### Splenocytes immune cell analysis

Flow cytometry was used to analyze immune cell populations in spleens collected from vaccinated C57BL/6J mice. On day 76, three weeks after the second boost, mice were euthanized and spleens were processed into single-cell suspensions, aliquoted, and stored frozen until further analysis, as previously described^36^.

Frozen aliquots were thawed in RPMI 1640 medium pre-warmed to 37°C, centrifuged, and resuspended in FACS buffer. Cells were stained for 30 minutes at 4°C with either B220-PerCP-Cy5.5 (Invitrogen), GL7-Alexa Fluor 647 (BioLegend), FAS(CD95)-PE-Cy7 (Miltenyi Biotec), CD3-APC-Cy7 (BioLegend), CD4-BV421 (BioLegend), and CD8a-FITC (BioLegend), or with B220-PerCP-Cy5.5, IgG-BV605 (BioLegend), IgM-FITC (BioLegend), and Mpox A28 conjugated to both Streptavidin-PE and Streptavidin-Alexa Fluor 647. Following a final wash, stained cells were analyzed by flow cytometry.

### Viral load

Lungs from infected mice were stored in PBS containing 0.1% BSA until homogenization using gentleMACS™ M Tubes and a gentleMACS™ Dissociator (Miltenyi Biotec). Viral load was quantified by titration on a confluent monolayer of Vero cells, as described above, and fluorescent foci were visualized and analyzed using the IncuCyte live-cell analysis system.

### X-ray crystallography

For structural studies, a synthetic gene encoding the head domain (amino acids 1–397) of A26 from the Vaccinia virus strain Western Reserve (Uniprot code: P24758), tagged at the C-terminus with a Strep-tag, was inserted into a pET-16b vector. The plasmid was transformed into *E. coli* BL21 (DE3) cells (New England Biolabs), and protein expression was induced overnight at 16°C with 0.5 mM isopropyl β-D-1-thiogalactopyranoside (IPTG). Cells from 3 liters of culture were harvested, resuspended in 40 mL of cold lysis buffer supplemented with one tablet of complete protease inhibitor (Roche), and lysed by sonication. Insoluble material was removed by centrifugation at 20,000 × g for 30 minutes at 4°C. The recombinant protein was purified by affinity chromatography using a 5 mL StrepTrap™ HP column (Cytiva), followed by size-exclusion chromatography in TN8 buffer (10 mM Tris-HCl pH 8, 100 mM NaCl) on a Superdex 200 column (Cytiva) to remove aggregates. Purity was assessed by SDS-PAGE and the final protein yield was 2 mg/L, as determined by NanoDrop spectrophotometer. To produce full-length antibodies, Expi293 cells were seeded at 3 × 10L cells/mL and transfected in 200 mL cultures with 200 μg each of heavy and light chain expression vectors using 200 μL of FectoPro transfection reagent. Cultures were incubated for 5 days at 37°C in FreeStyle medium. Supernatants were clarified by centrifugation at 4,000 rpm for 30 minutes at 4°C, followed by filtration through a 0.45 μm membrane. Antibodies were purified using a 5 mL Protein G column (Cytiva), eluted with 100 mM glycine pH 2.7, and further purified by size-exclusion chromatography in TN8 buffer on a Superdex 200 column. Purity was verified by SDS-PAGE, and concentrations were measured using NanoDrop. Final yields were approximately 30 mg/L for 10M2146 and 22 mg/L for 8M2110. Fab fragments of 8M2110 and 10M2146 were generated by papain digestion. For each antibody, 2.5 mg were incubated with 500 μL of 50% papain slurry (Thermo, cat. no. 20341). Fc fragments were removed using a Protein A column (Cytiva), and the monomeric Fab fraction was purified by size-exclusion chromatography.

For crystallization, purification tags were removed by overnight digestion with 1.5 units of thrombin (Cytiva) per 0.1 mg of protein at 4°C. Complexes were assembled by incubating A26 with a molar excess of Fab (8M2110 or 10M2146) and an anti-Fab VHH that binds the constant region of the light chain and stabilizes the hinge^76,77^. Complexes were purified by size-exclusion chromatography in TN8 buffer and concentrated to 3–5 mg/mL. Crystallization of the A26:8M2110:VHH complex was performed using 20% (w/v) PEG 3350 and 0.2 M ammonium citrate while the A26:10M2146:VHH complex was crystallized using 0.1 M MgCl₂, 0.1 M Tris-HCl pH 8.5, and 17% (w/v) PEG 20,000. In both cases, crystals were cryoprotected with ethylene glycol and diffraction data were collected at the SOLEIL synchrotron (St. Aubin, France) on beamlines PROXIMA-1 and PROXIMA-2. Diffraction data were processed using XDS (version January 10, 2022)^78^, and initial phases were obtained using PHASER software^79^ with the A26 model (PDB: 6A9S)^31^ as a search template. Structural models were built and refined iteratively using *phenix.refine* (Phenix version 1.19.2-4158)^80^ and *Coot^81^*, applying isotropic B-factor and TLS group refinement strategies. Model validation was performed with MolProbity^82^. Crystallographic statistics are provided. Coordinates and structure factors have been deposited in the Protein Data Bank under accession codes 9QT3 and 9QTA. Structural figures were generated with PyMOL v3.0.3 (Schrödinger, LLC).

### Biopanning and epitope prediction

Biopanning was performed as previously described^51^. Duplicate wells containing 2 μg/mL of anti-H3 mAb (7M1162) were incubated for 1 hour with protein-G magnetic beads (Thermo Scientific) in a blocking buffer containing TBS×1 and 3% BSA. The antibody-bead mixture was placed on a magnetic stand, and the supernatant containing unbound mAbs was discarded. The beads were then washed three times with washing buffer (TBS×1 and 0.05% Tween-20) to remove non-specifically bound proteins. Approximately 10¹¹ phages from the random-peptide library were added to the antibody-bound magnetic beads in blocking buffer and incubated for 1 hour. After incubation, the unbound phages were removed by discarding the supernatant, and the beads were washed three times with washing buffer. Affinity-selected phages were eluted using a low-pH solution (3 mg/mL BSA and 0.1 M glycine, pH 2.2) and immediately neutralized with 1 M Tris-HCl, pH 9.1. For phage amplification, *E. coli* DH5αF⁺ were infected with the affinity-selected phages and grown overnight at 37°C in liquid Terrific Broth (TB) and phosphate buffer (PB) medium containing 20 µg/mL tetracycline. The infected bacteria were then centrifuged, and the supernatant containing phages was incubated with 0.4 volumes of PEG-NaCl solution for 2 hours at 4°C. Following incubation, the mixture was centrifuged, the supernatant discarded, and the affinity-selected phages were resuspended in TBS×1. Each well underwent three rounds of Biopanning, with amplification steps between rounds.

Dot blot analysis was performed to confirm the binding of mAb 7M1162 to affinity-selected phages. *E. coli* DH5αF⁺ were infected with affinity-selected phages from the third round of biopanning and grown overnight at 37°C on Luria broth (LB) plates containing 20 µg/mL tetracycline (Holland Moran). Single bacterial colonies were individually picked and cultured overnight at 37°C in TB+PB medium containing 20 µg/mL tetracycline. The cultures were then centrifuged, and the supernatant containing phages was incubated with 0.4 volumes of PEG-NaCl solution for 2 hours at 4°C. After incubation, the mixture was centrifuged, the supernatant discarded, and affinity-selected phages were resuspended in TBS×1. Single-colony-derived phages were transferred onto nitrocellulose membranes (Tamar Ltd.) using negative pressure. Membranes were blocked for 2 hours at room temperature (RT) in a blocking solution containing TBS×1, 5% skim milk, and 0.05% Tween-20, followed by a single wash with TBS×1 and 0.05% Tween-20. Mab 7M1162 used for Biopanning was diluted to 2 μg/mL in blocking solution and incubated with the membranes for 1 hour at RT. Following three washes with washing buffer, membranes were incubated with an anti-human IgG HRP-conjugated secondary antibody (1:5000 dilution in blocking solution) for 45 minutes at RT. After five additional washes, ECL substrate was added, and Western blot images were acquired. Dot blot-positive phages were regrown, isolated, and revalidated.

Plasmids containing the DNA of validated phages were isolated and sent for Sanger sequencing to determine the peptide insert on the pVIII protein. Unique peptide sequences were used for epitope prediction based on the VACV H3 structure PDB:5EJ0^83^ using the PepSurf^51^ software. Amino acid sequences of mAb 7M1162 and full-length Mpox H3 were modeled together using AlphaFold3^52^ and analyzed using PyMOL software. Amino acid residues within 5 Å of 7M1162 in the predicted structure and forming a polar binding interface were inferred as possible binding sites.

## QUANTIFICATION AND STATISTICAL ANALYSIS

Statistical analysis was carried out using the GraphPad Prism software version 9.4.1. Analysis of Flow Cytometry data was carried out using FlowJo software version 10.8. Fluorescent cell images were analyzed using the IncuCyte live-cell analysis system for VACV and Harmony software for Mpox. Protein 3D structures were visualized using Pymol software version 3.1.3.1. Statistical details of each experiment can be found in the figure legend associated with each figure. Images were created by Biorender.

## AUTHOR CONTRIBUTIONS

NTF and RY conceived the project, planned and supervised the experiments and data analyses, and wrote the manuscript together with PGC and OS. NF collected human whole blood samples with assistance from DH and ES, and RY processed the samples. RY and NBS performed antigen production, single B cell sorting, monoclonal antibody cloning, and binding characterization experiments. RY and KR carried out VACV propagation with assistance from MRA, SS, and TK, under the supervision of OK. In vitro VACV and Mpox assays were performed by RY, KR, MH, FGB, FP, and JP. ZF provided critical feedback on the complement assays. In vivo experiments were conducted by RY, LA, and KR. LB performed all structural analyses, data collection, and figure preparation with assistance from PGC. RY, GO, and KP conducted Biopanning experiments and in silico modeling. RY and NR performed the TEM experiments under the supervision of LAA.

## ACKNOWLEDGMENTS

We thank Bernard Moss for providing us with the VACV strains: VACV-WR-vFire, VACV-IHDJ-vFire and VACV-Copenhagen-vFire. We thank Ohad Mazor, Tomer Israely and Hadas Tamir from the Israel Institute for Biological Research for their support. We thank Dr. Vered Holdengrerber for her assistance with TEM measurements. We thank Vice President of Research and Development of Tel Aviv University for funding. We thank Eli Gelman, Tel Aviv University’s Executive Council, and Ariel Porat, the President of Tel Aviv University for their support at the beginning of this project. We thank Israel Science Foundation (ISF) grants [3136/22] and [638/23] to NTF; Binational Science Foundation (BSF) [01031771] to NTF; BMGF INV-058519 to NTF. RY was supported by a PhD Scholarship from the Tel Aviv University Center for Combatting Pandemics. KP’s research is supported in part by a fellowship from the Edmond J. Safra Center for Bioinformatics at Tel Aviv University. NR thanks the Sagol Center for Regenerative Medicine for financial support. The OS lab is funded by Institut Pasteur, Fondation pour la Recherche Médicale (FRM), ANRS-MIE, the Vaccine Research Institute (VRI) (ANR-10-LABX-77), Labex IBEID (ANR-10-LABX-62-IBEID), the HERA projects DURABLE (grant 101102733) and LEAPS. Lastly, we thank all the donors who participated in this study.

## REFERENCES

1. Sklenovska, N., and Van Ranst, M. (2018). Emergence of Monkeypox as the Most Important Orthopoxvirus Infection in Humans. Front Public Health 6, 241. 10.3389/fpubh.2018.00241.

2. Thornhill, J.P., Gandhi, M., and Orkin, C. (2024). Mpox: The Reemergence of an Old Disease and Inequities. Annu Rev Med 75, 159–175. 10.1146/annurev-med-080122-030714.

3. McCollum, A.M., and Damon, I.K. (2014). Human monkeypox. Clin Infect Dis 58, 260–267. 10.1093/cid/cit703.

4. Lum, F.M., Torres-Ruesta, A., Tay, M.Z., Lin, R.T.P., Lye, D.C., Renia, L., and Ng, L.F.P. (2022). Monkeypox: disease epidemiology, host immunity and clinical interventions. Nat Rev Immunol 22, 597–613. 10.1038/s41577-022-00775-4.

5. Zebardast, A., Latifi, T., Shafiei-Jandaghi, N.Z., Gholami Barzoki, M., and Shatizadeh Malekshahi, S. (2023). Plausible reasons for the resurgence of Mpox (formerly Monkeypox): an overview. Trop Dis Travel Med Vaccines 9, 23. 10.1186/s40794-023-00209-6.

6. Reed, K.D., Melski, J.W., Graham, M.B., Regnery, R.L., Sotir, M.J., Wegner, M.V., Kazmierczak, J.J., Stratman, E.J., Li, Y., Fairley, J.A., et al. (2004). The detection of monkeypox in humans in the Western Hemisphere. N Engl J Med 350, 342–350. 10.1056/NEJMoa032299.

7. Organization, W.H. (2024). Mpox: Multi-country External Situation Report n.42.

8. McQuiston, J.H., Braden, C.R., Bowen, M.D., McCollum, A.M., McDonald, R., Carnes, N., Carter, R.J., Christie, A., Doty, J.B., Ellington, S., et al. (2023). The CDC Domestic Mpox Response - United States, 2022-2023. MMWR Morb Mortal Wkly Rep 72, 547-552. 10.15585/mmwr.mm7220a2.

9. Zaeck, L.M., Lamers, M.M., Verstrepen, B.E., Bestebroer, T.M., van Royen, M.E., Gotz, H., Shamier, M.C., van Leeuwen, L.P.M., Schmitz, K.S., Alblas, K., et al. (2023). Low levels of monkeypox virus-neutralizing antibodies after MVA-BN vaccination in healthy individuals. Nat Med 29, 270–278. 10.1038/s41591-022-02090-w.

10. Collier, A.Y., McMahan, K., Jacob-Dolan, C., Liu, J., Borducchi, E.N., Moss, B., and Barouch, D.H. (2024). Decline of Mpox Antibody Responses After Modified Vaccinia Ankara-Bavarian Nordic Vaccination. JAMA 332, 1669–1672. 10.1001/jama.2024.20951.

11. Priyamvada, L., Carson, W.C., Ortega, E., Navarra, T., Tran, S., Smith, T.G., Pukuta, E., Muyamuna, E., Kabamba, J., Nguete, B.U., et al. (2022). Serological responses to the MVA-based JYNNEOS monkeypox vaccine in a cohort of participants from the Democratic Republic of Congo. Vaccine 40, 7321–7327. 10.1016/j.vaccine.2022.10.078.

12. Deputy, N.P., Deckert, J., Chard, A.N., Sandberg, N., Moulia, D.L., Barkley, E., Dalton, A.F., Sweet, C., Cohn, A.C., Little, D.R., et al. (2023). Vaccine Effectiveness of JYNNEOS against Mpox Disease in the United States. N Engl J Med 388, 2434–2443. 10.1056/NEJMoa2215201.

13. Pischel, L., Martini, B.A., Yu, N., Cacesse, D., Tracy, M., Kharbanda, K., Ahmed, N., Patel, K.M., Grimshaw, A.A., Malik, A.A., et al. (2024). Vaccine effectiveness of 3rd generation mpox vaccines against mpox and disease severity: A systematic review and meta-analysis. Vaccine 42, 126053. 10.1016/j.vaccine.2024.06.021.

14. Crandell, J., Monteiro, V.S., Pischel, L., Fang, Z., Conde, L., Zhong, Y., Lawres, L., de Asis, G.M., Maciel, G., Zaleski, A., et al. (2025). The impact of orthopoxvirus vaccination and Mpox infection on cross-protective immunity: a multicohort observational study. Lancet Microbe, 101098. 10.1016/j.lanmic.2025.101098.

15. Cohn, H., Bloom, N., Cai, G.Y., Clark, J.J., Tarke, A., Bermudez-Gonzalez, M.C., Altman, D.R., Lugo, L.A., Lobo, F.P., Marquez, S., et al. (2023). Mpox vaccine and infection-driven human immune signatures: an immunological analysis of an observational study. Lancet Infect Dis 23, 1302–1312. 10.1016/S1473-3099(23)00352-3.

16. Otter, A.D., Jones, S., Hicks, B., Bailey, D., Callaby, H., Houlihan, C., Rampling, T., Gordon, N.C., Selman, H., Satheshkumar, P.S., et al. (2023). Monkeypox virus-infected individuals mount comparable humoral immune responses as Smallpox-vaccinated individuals. Nat Commun 14, 5948. 10.1038/s41467-023-41587-x.

17. Yefet, R., Friedel, N., Tamir, H., Polonsky, K., Mor, M., Cherry-Mimran, L., Taleb, E., Hagin, D., Sprecher, E., Israely, T., and Freund, N.T. (2023). Monkeypox infection elicits strong antibody and B cell response against A35R and H3L antigens. iScience 26, 105957. 10.1016/j.isci.2023.105957.

18. Moss, B. (2012). Poxvirus cell entry: how many proteins does it take? Viruses 4, 688–707. 10.3390/v4050688.

19. Smith, G.L., Vanderplasschen, A., and Law, M. (2002). The formation and function of extracellular enveloped vaccinia virus. J Gen Virol 83, 2915–2931. 10.1099/0022-1317-83-12-2915.

20. Blasco, R., and Moss, B. (1992). Role of cell-associated enveloped vaccinia virus in cell-to-cell spread. J Virol 66, 4170–4179. 10.1128/JVI.66.7.4170-4179.1992.

21. Gilchuk, I., Gilchuk, P., Sapparapu, G., Lampley, R., Singh, V., Kose, N., Blum, D.L., Hughes, L.J., Satheshkumar, P.S., Townsend, M.B., et al. (2016). Cross-Neutralizing and Protective Human Antibody Specificities to Poxvirus Infections. Cell 167, 684–694 e689. 10.1016/j.cell.2016.09.049.

22. Liu, J., Wang, X., Zhang, Y., Liu, C., Zhang, M., Li, C., Liu, P., Li, S., Wei, K., Cai, Y., et al. (2025). Immunogenicity of monkeypox virus surface proteins and cross-reactive antibody responses in vaccinated and infected individuals: implications for vaccine and therapeutic development. Infect Dis Poverty 14, 12. 10.1186/s40249-025-01280-1.

23. Riccardo, V., and Pablo, G.C. (2023). Neutralization Determinants on Poxviruses. Viruses 15. 10.3390/v15122396.

24. Hou, R., Jiang, Q., Cheng, M., Dai, J., Yang, H., Yuan, J., Li, X., Tang, X., and Yu, H. (2024). Identification of neutralizing antibodies against monkeypox virus using high-throughput sequencing of A35(+)H3L(+)B cells from patients with convalescent monkeypox. Virus Res 347, 199437. 10.1016/j.virusres.2024.199437.

25. Zhao, R., Wu, L., Sun, J., Liu, D., Han, P., Gao, Y., Zhang, Y., Xu, Y., Qu, X., Wang, H., et al. (2024). Two noncompeting human neutralizing antibodies targeting MPXV B6 show protective effects against orthopoxvirus infections. Nat Commun 15, 4660. 10.1038/s41467-024-48312-2.

26. Noy-Porat, T., Tamir, H., Alcalay, R., Rosenfeld, R., Epstein, E., Cherry, L., Achdout, H., Erez, N., Politi, B., Yahalom-Ronen, Y., et al. (2023). Generation of recombinant mAbs to vaccinia virus displaying high affinity and potent neutralization. Microbiol Spectr 11, e0159823. 10.1128/spectrum.01598-23.

27. Tamir, H., Noy-Porat, T., Melamed, S., Cherry-Mimran, L., Barlev-Gross, M., Alcalay, R., Yahalom-Ronen, Y., Achdout, H., Politi, B., Erez, N., et al. (2024). Synergistic effect of two human-like monoclonal antibodies confers protection against orthopoxvirus infection. Nat Commun 15, 3265. 10.1038/s41467-024-47328-y.

28. McCausland, M.M., Benhnia, M.R., Crickard, L., Laudenslager, J., Granger, S.W., Tahara, T., Kubo, R., Koriazova, L., Kato, S., and Crotty, S. (2010). Combination therapy of vaccinia virus infection with human anti-H3 and anti-B5 monoclonal antibodies in a small animal model. Antivir Ther 15, 661–675. 10.3851/IMP1573.

29. Tai, W., Tian, C., Shi, H., Chai, B., Yu, X., Zhuang, X., Dong, P., Li, M., Yin, Q., Feng, S., et al. (2025). An mRNA vaccine against monkeypox virus inhibits infection by co-activation of humoral and cellular immune responses. Nat Commun 16, 2971. 10.1038/s41467-025-58328-x.

30. Pugh, C., Keasey, S., Korman, L., Pittman, P.R., and Ulrich, R.G. (2014). Human antibody responses to the polyclonal Dryvax vaccine for smallpox prevention can be distinguished from responses to the monoclonal replacement vaccine ACAM2000. Clin Vaccine Immunol 21, 877–885. 10.1128/CVI.00035-14.

31. Chang, H.W., Yang, C.H., Luo, Y.C., Su, B.G., Cheng, H.Y., Tung, S.Y., Carillo, K.J.D., Liao, Y.T., Tzou, D.M., Wang, H.C., and Chang, W. (2019). Vaccinia viral A26 protein is a fusion suppressor of mature virus and triggers membrane fusion through conformational change at low pH. PLoS Pathog 15, e1007826. 10.1371/journal.ppat.1007826.

32. Chang, S.J., Chang, Y.X., Izmailyan, R., Tang, Y.L., and Chang, W. (2010). Vaccinia virus A25 and A26 proteins are fusion suppressors for mature virions and determine strain-specific virus entry pathways into HeLa, CHO-K1, and L cells. J Virol 84, 8422-8432. 10.1128/JVI.00599-10.

33. Mor, M., Werbner, M., Alter, J., Safra, M., Chomsky, E., Lee, J.C., Hada-Neeman, S., Polonsky, K., Nowell, C.J., Clark, A.E., et al. (2021). Multi-clonal SARS-CoV-2 neutralization by antibodies isolated from severe COVID-19 convalescent donors. PLoS Pathog 17, e1009165. 10.1371/journal.ppat.1009165.

34. Watson, A., Li, H., Ma, B., Weiss, R., Bendayan, D., Abramovitz, L., Ben-Shalom, N., Mor, M., Pinko, E., Bar Oz, M., et al. (2021). Human antibodies targeting a Mycobacterium transporter protein mediate protection against tuberculosis. Nat Commun 12, 602. 10.1038/s41467-021-20930-0.

35. Wardemann, H., Yurasov, S., Schaefer, A., Young, J.W., Meffre, E., and Nussenzweig, M.C. (2003). Predominant autoantibody production by early human B cell precursors. Science 301, 1374–1377. 10.1126/science.1086907.

36. Ben-Shalom, N., Sandbank, E., Abramovitz, L., Hezroni, H., Levine, T., Trachtenberg, E., Fogel, N., Mor, M., Yefet, R., Stoler-Barak, L., et al. (2023). beta2-adrenergic signaling promotes higher-affinity B cells and antibodies. Brain Behav Immun 113, 66–82. 10.1016/j.bbi.2023.06.020.

37. Imani, S., Aminnezhad, S., Alikarami, M., Abedi, Z., Mosleh, I.S., Maghsoudloo, M., and Taheri, Z. (2024). Exploration of drug repurposing for Mpox outbreaks targeting gene signatures and host-pathogen interactions. Sci Rep 14, 29436. 10.1038/s41598-024-79897-9.

38. Ahmed, S.F., Sohail, M.S., Quadeer, A.A., and McKay, M.R. (2022). Vaccinia-Virus-Based Vaccines Are Expected to Elicit Highly Cross-Reactive Immunity to the 2022 Monkeypox Virus. Viruses 14. 10.3390/v14091960.

39. Sok, D., Briney, B., Jardine, J.G., Kulp, D.W., Menis, S., Pauthner, M., Wood, A., Lee, E.C., Le, K.M., Jones, M., et al. (2016). Priming HIV-1 broadly neutralizing antibody precursors in human Ig loci transgenic mice. Science 353, 1557–1560. 10.1126/science.aah3945.

40. Freund, N.T., Horwitz, J.A., Nogueira, L., Sievers, S.A., Scharf, L., Scheid, J.F., Gazumyan, A., Liu, C., Velinzon, K., Goldenthal, A., et al. (2015). A New Glycan-Dependent CD4-Binding Site Neutralizing Antibody Exerts Pressure on HIV-1 In Vivo. PLoS Pathog 11, e1005238. 10.1371/journal.ppat.1005238.

41. Scharf, L., West, A.P., Jr., Gao, H., Lee, T., Scheid, J.F., Nussenzweig, M.C., Bjorkman, P.J., and Diskin, R. (2013). Structural basis for HIV-1 gp120 recognition by a germ-line version of a broadly neutralizing antibody. Proc Natl Acad Sci U S A 110, 6049–6054. 10.1073/pnas.1303682110.

42. Hubert, M., Guivel-Benhassine, F., Bruel, T., Porrot, F., Planas, D., Vanhomwegen, J., Wiedemann, A., Burrel, S., Marot, S., Palich, R., et al. (2023). Complement-dependent mpox-virus-neutralizing antibodies in infected and vaccinated individuals. Cell Host Microbe 31, 937–948 e934. 10.1016/j.chom.2023.05.001.

43. Bengali, Z., Satheshkumar, P.S., and Moss, B. (2012). Orthopoxvirus species and strain differences in cell entry. Virology 433, 506–512. 10.1016/j.virol.2012.08.044.

44. Monzon, S., Varona, S., Negredo, A., Vidal-Freire, S., Patino-Galindo, J.A., Ferressini-Gerpe, N., Zaballos, A., Orviz, E., Ayerdi, O., Munoz-Gomez, A., et al. (2024). Monkeypox virus genomic accordion strategies. Nat Commun 15, 3059. 10.1038/s41467-024-46949-7.

45. Senkevich, T.G., Yutin, N., Wolf, Y.I., Koonin, E.V., and Moss, B. (2021). Ancient Gene Capture and Recent Gene Loss Shape the Evolution of Orthopoxvirus-Host Interaction Genes. mBio 12, e0149521. 10.1128/mBio.01495-21.

46. Mendoza, P., Gruell, H., Nogueira, L., Pai, J.A., Butler, A.L., Millard, K., Lehmann, C., Suarez, I., Oliveira, T.Y., Lorenzi, J.C.C., et al. (2018). Combination therapy with anti-HIV-1 antibodies maintains viral suppression. Nature 561, 479–484. 10.1038/s41586-018-0531-2.

47. Howard, A.R., Senkevich, T.G., and Moss, B. (2008). Vaccinia virus A26 and A27 proteins form a stable complex tethered to mature virions by association with the A17 transmembrane protein. J Virol 82, 12384–12391. 10.1128/JVI.01524-08.

48. Bublil, E.M., Freund, N.T., Mayrose, I., Penn, O., Roitburd-Berman, A., Rubinstein, N.D., Pupko, T., and Gershoni, J.M. (2007). Stepwise prediction of conformational discontinuous B-cell epitopes using the Mapitope algorithm. Proteins 68, 294–304. 10.1002/prot.21387.

49. Gershoni, J.M., Roitburd-Berman, A., Siman-Tov, D.D., Tarnovitski Freund, N., and Weiss, Y. (2007). Epitope mapping: the first step in developing epitope-based vaccines. BioDrugs 21, 145–156. 10.2165/00063030-200721030-00002.

50. Mayrose, I., Penn, O., Erez, E., Rubinstein, N.D., Shlomi, T., Freund, N.T., Bublil, E.M., Ruppin, E., Sharan, R., Gershoni, J.M., et al. (2007). Pepitope: epitope mapping from affinity-selected peptides. Bioinformatics 23, 3244–3246. 10.1093/bioinformatics/btm493.

51. Mayrose, I., Shlomi, T., Rubinstein, N.D., Gershoni, J.M., Ruppin, E., Sharan, R., and Pupko, T. (2007). Epitope mapping using combinatorial phage-display libraries: a graph-based algorithm. Nucleic Acids Res 35, 69–78. 10.1093/nar/gkl975.

52. Abramson, J., Adler, J., Dunger, J., Evans, R., Green, T., Pritzel, A., Ronneberger, O., Willmore, L., Ballard, A.J., Bambrick, J., et al. (2024). Addendum: Accurate structure prediction of biomolecular interactions with AlphaFold 3. Nature 636, E4. 10.1038/s41586-024-08416-7.

53. Benhnia, M.R., Maybeno, M., Blum, D., Aguilar-Sino, R., Matho, M., Meng, X., Head, S., Felgner, P.L., Zajonc, D.M., Koriazova, L., et al. (2013). Unusual features of vaccinia virus extracellular virion form neutralization resistance revealed in human antibody responses to the smallpox vaccine. J Virol 87, 1569–1585. 10.1128/JVI.02152-12.

54. Benhnia, M.R., McCausland, M.M., Moyron, J., Laudenslager, J., Granger, S., Rickert, S., Koriazova, L., Kubo, R., Kato, S., and Crotty, S. (2009). Vaccinia virus extracellular enveloped virion neutralization in vitro and protection in vivo depend on complement. J Virol 83, 1201–1215. 10.1128/JVI.01797-08.

55. Cohen, M.E., Xiao, Y., Eisenberg, R.J., Cohen, G.H., and Isaacs, S.N. (2011). Antibody against extracellular vaccinia virus (EV) protects mice through complement and Fc receptors. PLoS One 6, e20597. 10.1371/journal.pone.0020597.

56. Holley, J., Sumner, R.P., Lant, S., Ribeca, P., Ulaeto, D., and Maluquer de Motes, C. (2021). Engineered Promoter-Switched Viruses Reveal the Role of Poxvirus Maturation Protein A26 as a Negative Regulator of Viral Spread. J Virol 95, e0101221. 10.1128/JVI.01012-21.

57. Burton, D.R. (2023). Antiviral neutralizing antibodies: from in vitro to in vivo activity. Nat Rev Immunol 23, 720–734. 10.1038/s41577-023-00858-w.

58. Freund, N.T., Wang, H., Scharf, L., Nogueira, L., Horwitz, J.A., Bar-On, Y., Golijanin, J., Sievers, S.A., Sok, D., Cai, H., et al. (2017). Coexistence of potent HIV-1 broadly neutralizing antibodies and antibody-sensitive viruses in a viremic controller. Sci Transl Med 9. 10.1126/scitranslmed.aal2144.

59. Suryadevara, N., Shrihari, S., Gilchuk, P., VanBlargan, L.A., Binshtein, E., Zost, S.J., Nargi, R.S., Sutton, R.E., Winkler, E.S., Chen, E.C., et al. (2021). Neutralizing and protective human monoclonal antibodies recognizing the N-terminal domain of the SARS-CoV-2 spike protein. Cell 184, 2316–2331 e2315. 10.1016/j.cell.2021.03.029.

60. Wrammert, J., Smith, K., Miller, J., Langley, W.A., Kokko, K., Larsen, C., Zheng, N.Y., Mays, I., Garman, L., Helms, C., et al. (2008). Rapid cloning of high-affinity human monoclonal antibodies against influenza virus. Nature 453, 667–671. 10.1038/nature06890.

61. Strassburg, M.A. (1982). The global eradication of smallpox. Am J Infect Control 10, 53–59. 10.1016/0196-6553(82)90003-7.

62. Volz, A., and Sutter, G. (2017). Modified Vaccinia Virus Ankara: History, Value in Basic Research, and Current Perspectives for Vaccine Development. Adv Virus Res 97, 187–243. 10.1016/bs.aivir.2016.07.001.

63. Freyn, A.W., Atyeo, C., Earl, P.L., Americo, J.L., Chuang, G.Y., Natarajan, H., Frey, T.R., Gall, J.G., Moliva, J.I., Hunegnaw, R., et al. (2023). An mpox virus mRNA-lipid nanoparticle vaccine confers protection against lethal orthopoxviral challenge. Sci Transl Med 15, eadg3540. 10.1126/scitranslmed.adg3540.

64. Mucker, E.M., Freyn, A.W., Bixler, S.L., Cizmeci, D., Atyeo, C., Earl, P.L., Natarajan, H., Santos, G., Frey, T.R., Levin, R.H., et al. (2024). Comparison of protection against mpox following mRNA or modified vaccinia Ankara vaccination in nonhuman primates. Cell 187, 5540–5553 e5510. 10.1016/j.cell.2024.08.043.

65. Wang, H., Yin, P., Zheng, T., Qin, L., Li, S., Han, P., Qu, X., Wen, J., Ding, H., Wu, J., et al. (2024). Rational design of a ‘two-in-one’ immunogen DAM drives potent immune response against mpox virus. Nat Immunol 25, 307–315. 10.1038/s41590-023-01715-7.

66. Zuiani, A., Dulberger, C.L., De Silva, N.S., Marquette, M., Lu, Y.J., Palowitch, G.M., Dokic, A., Sanchez-Velazquez, R., Schlatterer, K., Sarkar, S., et al. (2024). A multivalent mRNA monkeypox virus vaccine (BNT166) protects mice and macaques from orthopoxvirus disease. Cell 187, 1363–1373 e1312. 10.1016/j.cell.2024.01.017.

67. Jacob-Dolan, C., Ty, D., Hope, D., McMahan, K., Liu, J., Powers, O.C., Cotter, C.A., Sciacca, M., Wu, C., Borducchi, E., et al. (2024). Comparison of the immunogenicity and protective efficacy of ACAM2000, MVA, and vectored subunit vaccines for Mpox in rhesus macaques. Sci Transl Med 16, eadl4317. 10.1126/scitranslmed.adl4317.

68. Ye, Q., Zhang, D., Zhang, R.R., Xu, Q., Huang, X.Y., Huang, B., Sun, M.X., Cong, Z., Zhu, L., Ma, J., et al. (2024). A penta-component mpox mRNA vaccine induces protective immunity in nonhuman primates. Nat Commun 15, 10611. 10.1038/s41467-024-54909-4.

69. Baliere, C., Hourdel, V., Kwasiborski, A., Grassin, Q., Feher, M., Hoinard, D., Vanhomwegen, J., Taieb, F., Consigny, P.H., Manuguerra, J.C., et al. (2023). Complete Genome Sequences of Monkeypox Virus from a French Clinical Sample and the Corresponding Isolated Strain, Obtained Using Nanopore Sequencing. Microbiol Resour Announc 12, e0000923. 10.1128/mra.00009-23.

70. Cohen-Gihon, I., Israeli, O., Shifman, O., Erez, N., Melamed, S., Paran, N., Beth-Din, A., and Zvi, A. (2020). Identification and Whole-Genome Sequencing of a Monkeypox Virus Strain Isolated in Israel. Microbiol Resour Announc 9. 10.1128/MRA.01524-19.

71. Qin, L., Favis, N., Famulski, J., and Evans, D.H. (2015). Evolution of and evolutionary relationships between extant vaccinia virus strains. J Virol 89, 1809–1824. 10.1128/JVI.02797-14.

72. Tiller, T., Meffre, E., Yurasov, S., Tsuiji, M., Nussenzweig, M.C., and Wardemann, H. (2008). Efficient generation of monoclonal antibodies from single human B cells by single cell RT-PCR and expression vector cloning. J Immunol Methods 329, 112–124. 10.1016/j.jim.2007.09.017.

73. Wolffe, E.J., Vijaya, S., and Moss, B. (1995). A myristylated membrane protein encoded by the vaccinia virus L1R open reading frame is the target of potent neutralizing monoclonal antibodies. Virology 211, 53–63. 10.1006/viro.1995.1378.

74. Su, H.P., Golden, J.W., Gittis, A.G., Hooper, J.W., and Garboczi, D.N. (2007). Structural basis for the binding of the neutralizing antibody, 7D11, to the poxvirus L1 protein. Virology 368, 331-341. 10.1016/j.virol.2007.06.042.

75. Postal, J., Guivel-Benhassine, F., Porrot, F., Grassin, Q., Crook, J.M., Vernuccio, R., Caro, V., Vanhomwegen, J., Guardado-Calvo, P., Simon-Loriere, E., et al. (2025). Antiviral activity of tecovirimat against monkeypox virus clades 1a, 1b, 2a, and 2b. Lancet Infect Dis 25, e126-e127. 10.1016/S1473-3099(25)00014-3.

76. Ereno-Orbea, J., Sicard, T., Cui, H., Carson, J., Hermans, P., and Julien, J.P. (2018). Structural Basis of Enhanced Crystallizability Induced by a Molecular Chaperone for Antibody Antigen-Binding Fragments. J Mol Biol 430, 322–336. 10.1016/j.jmb.2017.12.010.

77. Bloch, J.S., Mukherjee, S., Kowal, J., Filippova, E.V., Niederer, M., Pardon, E., Steyaert, J., Kossiakoff, A.A., and Locher, K.P. (2021). Development of a universal nanobody-binding Fab module for fiducial-assisted cryo-EM studies of membrane proteins. Proc Natl Acad Sci U S A 118. 10.1073/pnas.2115435118.

78. Kabsch, W. (2010). Integration, scaling, space-group assignment and post-refinement. Acta Crystallogr D Biol Crystallogr 66, 133–144. 10.1107/S0907444909047374.

79. McCoy, A.J., Grosse-Kunstleve, R.W., Adams, P.D., Winn, M.D., Storoni, L.C., and Read, R.J. (2007). Phaser crystallographic software. J Appl Crystallogr 40, 658–674. 10.1107/S0021889807021206.

80. Afonine, P.V., Grosse-Kunstleve, R.W., Echols, N., Headd, J.J., Moriarty, N.W., Mustyakimov, M., Terwilliger, T.C., Urzhumtsev, A., Zwart, P.H., and Adams, P.D. (2012). Towards automated crystallographic structure refinement with phenix.refine. Acta Crystallogr D Biol Crystallogr 68, 352–367. 10.1107/S0907444912001308.

81. Emsley, P., Lohkamp, B., Scott, W.G., and Cowtan, K. (2010). Features and development of Coot. Acta Crystallogr D Biol Crystallogr 66, 486–501. 10.1107/S0907444910007493.

82. Williams, C.J., Headd, J.J., Moriarty, N.W., Prisant, M.G., Videau, L.L., Deis, L.N., Verma, V., Keedy, D.A., Hintze, B.J., Chen, V.B., et al. (2018). MolProbity: More and better reference data for improved all-atom structure validation. Protein Sci 27, 293–315. 10.1002/pro.3330.

83. Singh, K., Gittis, A.G., Gitti, R.K., Ostazeski, S.A., Su, H.P., and Garboczi, D.N. (2016). The Vaccinia Virus H3 Envelope Protein, a Major Target of Neutralizing Antibodies, Exhibits a Glycosyltransferase Fold and Binds UDP-Glucose. J Virol 90, 5020–5030. 10.1128/JVI.02933-15.

